# Transcriptional kinetic synergy: a complex landscape revealed by integrating modelling and synthetic biology

**DOI:** 10.1101/2020.08.31.276261

**Authors:** Rosa Martinez-Corral, Minhee Park, Kelly Biette, Dhana Friedrich, Clarissa Scholes, Ahmad S. Khalil, Jeremy Gunawardena, Angela H. DePace

## Abstract

Gene regulation involves synergistic interactions between transcription factors (TFs). Classical thermodynamic models offer a biophysical understanding of synergy based on binding cooperativity and regulated recruitment of RNA polymerase. However, transcription requires polymerase to transition through multiple states. Accordingly, recent work has suggested that ”kinetic synergy” can arise through TFs differentially regulating distinct steps of the transcription cycle. Disentangling both sources of synergy has been challenging. Here, we combine theory and experiment to analyze TFs binding to a single shared site, thereby removing simultaneous specific DNA binding. Using the graph-based linear framework, we integrate TF binding with regulation of the transcription cycle, and reveal a complex kinetic synergy landscape dependent on TF concentration, DNA binding and transcriptional activity. We exploit synthetic zinc-finger TF fusions to experimentally interrogate these predictions. Our results confirm that transcription cycle regulation must be integrated with recruitment for a quantitative understanding of transcriptional control.

## 3 Introduction

The regulation of transcription is a finely controlled process central to biology, biomedicine and bioengineering applications. At its core are transcription factors (TFs), proteins that bind specific sites on the DNA and directly or indirectly modulate the binding and activity of the RNA polymerase complex. In eukaryotes, multiple TFs, of the same and distinct types, collaborate to drive transcription through binding to gene regulatory regions called enhancers and promoters (Field and Adelman, 2020). Such ”combinatorial control” enables binding and response specificity (Wunderlich and Mirny, 2009; Georges et al., 2010), and expands the regulatory capabilities of the finite set of TFs encoded by an organism. A wealth of studies have characterised TF binding sites and binding profiles in model genes, genomes and random sequences (e.g. Smith et al., 2013; Vandel et al., 2019; Inukai et al., 2017). In turn, a long-standing goal of biomedicine and synthetic biology has been to exploit this type of information to anticipate the effect of mutations on cell regulation, to develop new and more refined pharmacological interventions, and to design next-generation synthetic circuits with more precise and robust functions. However, this is still a difficult task, in part because of the non-independent effects of the TFs that control a given gene (Ouyang et al., 2009; de Boer et al., 2020; Reiter et al., 2017; King et al., 2020; Nie et al., 2020).

When TFs interact to regulate transcription, the response to a combination of TFs is often not simply predicted by the responses to each of the TFs alone. Some models indicate that in the absence of interactions between TFs or sites, their combined effect should just be the addition of the individual outputs, and ”synergy” has been used to refer to deviation from this additive expectation (Carey et al., 1990; Herschlag and Johnson, 1993; Scholes et al., 2017). Under other models, ”synergy” is manifested as multiplicativity in the response (Bintu et al., 2005a). Alternatively, the term ”synergy” has been used to refer to nonlinear response to increasing TF concentrations (Carey, 1998), binding cooperativity (below), or a special form of it (Veitia, 2003; Michida et al., 2020). Here we use the term ”synergy” to refer to an increase in the expression output under two TFs in comparison to their individual effects, quantified by a functional, model-agnostic measure proposed in the Results section.

Synergy has commonly been understood through the lens of recruitment models of transcription, where the role of TFs is to regulate the binding of the RNA polymerase to the gene (Ptashne, 2005). Thermodynamic models of gene regulation offer a biophysical grounding for this view (Ackers et al., 1982; Bintu et al., 2005b,a). These models assume that TFs and polymerase bind to the DNA under thermodynamic equilibrium conditions. The free energy of each state determines its steady-state probability according to the Boltzmann distribution, and the transcription rate is treated as a function of the states of binding of the system. Synergy then emerges from direct or indirect cooperative binding interactions, where TFs enhance or reduce each other’s binding and that of the RNA polymerase to the DNA (e.g. Vashee et al., 1998; Ambrosetti et al., 2000; Spitz and Furlong, 2012; Frank et al., 2012; Goldstein et al., 2017; Estrada et al., 2016). Mechanistically, this can result from direct protein-protein interactions between adjacently-bound molecules, indirect interactions through a shared molecule or complex like Mediator (Carey et al., 1990; Malik and Roeder, 2010; Bashor et al., 2019) or through allosteric mechanisms (Biddle et al., 2021) mediated by nucleosomes (Mirny, 2010) or by DNA (Narasimhan et al., 2015).

Beyond recruitment of RNA polymerase to the gene, it is well known that eukaryotic transcription is a multi-step process that is tightly regulated at many points. Accordingly, it has been suggested that transcriptional regulation should be understood in terms of a transcription cycle (Fuda et al., 2009), involving for example the displacement of nucleosomes at the start site, post-translational modification of histones (Mao et al., 2010; Hansen and O'Shea, 2013; Cui et al., 2020), assembly of the transcriptional machinery, and post-translational modifications that regulate its activity and elongation rate (Jonkers and Lis, 2015; Core and Adelman, 2019). In agreement with this view, RNA polymerase has been found to be already bound on many inactive genes, suggesting that under certain scenarios activation does not rely on regulating polymerase recruitment, but modulating a subsequent step (Oven et al., 2007). Besides moving the focus away from the recruitment of the RNA polymerase, this view also implies non-equilibrium behaviour, given that ATP-dependent nucleosome remodelling and post-translational modifications involve energy dissipation. In this case, the steady-state behaviour of the system is determined by the individual rates of the various transitions. This is in contrast to the equilibrium situation of thermodynamic models, where only the ratios between the forward and backward rates matter for determining the steady-state of the system (Wong and Gunawardena, 2020).

Under this kinetic view, the possibility of ”kinetic synergy” was theoretically proposed. Imagine the simplest case where transcription is regulated by two steps, and two TFs have different biochemical functions such that one TF can preferentially enhance one step and the other TF can preferentially enhance the other. Then, when the two TFs are present together they can enhance each other’s effect and thus generate synergy (Herschlag and Johnson, 1993; Scholes et al., 2017). Importantly, this would enable synergy to emerge even in the absence of cooperative binding between TFs on the DNA; the TFs would not even need to be simultaneously present at the regulatory site.

Multiple lines of evidence make kinetic synergy very plausible. First, experimental work has shown that transcriptional activators can increase gene expression by different mechanisms. Blau et al. (1996) found that TF activation domains can either stimulate transcription initiation, elongation, or both, and more recent studies have continued to reveal that TFs use diverse mechanisms to regulate transcription and affect distinct steps of the transcription cycle (e.g. Fu et al., 2004; Rahl et al., 2010; Baluapuri et al., 2019). Along the same lines, Danko et al. (2013) reported differences in RNA polymerase II pausing depending upon treatment with E2 or TNF-alpha signals, which were attributed to the TFs downstream (ERα and NF-κB) acting on different transitions that regulate their target genes. Moreover, comparisons between regulation driven by homogeneous or heterogeneous sets of TFs have shown that heterogeneous sets often drive higher expression levels (Smith et al., 2013; Vanhille et al., 2015; Singh et al., 2021). In line with this, Keung et al. (2014) found evidence of synergistic activation between the viral activator VP16 and selected chromatin regulators in a reporter system. Similarly, the activity of many *Drosophila* TFs and cofactors was found to be highly context-dependent (Stampfel et al., 2015), suggesting that activation may require a particular combination of biochemical mechanisms.

Despite these observations, it is experimentally challenging to assess kinetic synergy given the difficulty of disentangling it from cooperative DNA-binding interactions between TFs. On the theoretical side, there has been a lack of tools to reason about kinetic synergy on biophysical grounds. As a first step, a recent theoretical study by our group showed that in a similar way to binding cooperativity, kinetic synergy can implement logical and analog computations (Scholes et al., 2017), and that it can generate a wide diversity of input/output relationships. However, in a similar way to other modelling work that considers transcription as a multi-step process (e.g Suter et al., 2011; Hansen and O'Shea, 2013; Rybakova et al., 2015), that model did not explicitly account for TF binding, and instead represented it indirectly through the effect of the TFs on the transition rates of the system. To our knowledge, there have been few attempts to explicitly model the interplay between TF binding, polymerase recruitment, and progression over the transcription cycle. Li et al. (2018) proposed a model that explicitly incorporated binding and transitions over the cycle, but assumed a time-scale separation between TF binding and the rest of the processes, with quasi-equilibrium in TF binding. However, both TF residence times and the half-life of certain biochemical steps in the transcription cycle may occur on similar timescales, on the order of several seconds or a few minutes (Methods, section 6.2), calling for more general models that bring together the binding-centered view of recruitment models with the regulation of the transcription cycle.

Here we exploit the graph-based linear framework (below) to propose a model of transcriptional control that explicitly accounts for TF binding and the regulation of polymerase recruitment, as well as the progression over the transcription cycle. In order to disentangle kinetic synergy from binding cooperativity, we focus on the emergence of synergy between TFs binding to a single, shared site. This scenario eliminates the possibility of TFs simultaneously bound to the DNA, thus removing cooperative binding between TFs. Experimentally, we build this system using engineered TFs where activation domains of a set of functionally diverse mammalian TFs are fused to a computationally designed zinc-finger (ZF) DNA binding domain predicted to bind only to an artificial site upstream of a reporter (Figure 1A) (Khalil et al., 2012; Keung et al., 2014; Park et al., 2019b; Israni et al., 2021). We propose a comprehensive measure of synergy where we compare the expression output when both TFs are present, to that when only one of them is present. By exploring the synergistic behaviour of the model in parameter space, we find that a diversity of behaviors can emerge in this setup, for which we find experimental evidence. Our model reveals a complex synergy landscape, shaped by the interplay between the activation effect of the TFs and their binding kinetics. This highlights the relevance of considering genomic context, binding and biochemical function together when characterizing TFs, and illuminates how functional interactions between TFs may contribute to eukaryotic transcriptional control.

**Figure 1:**
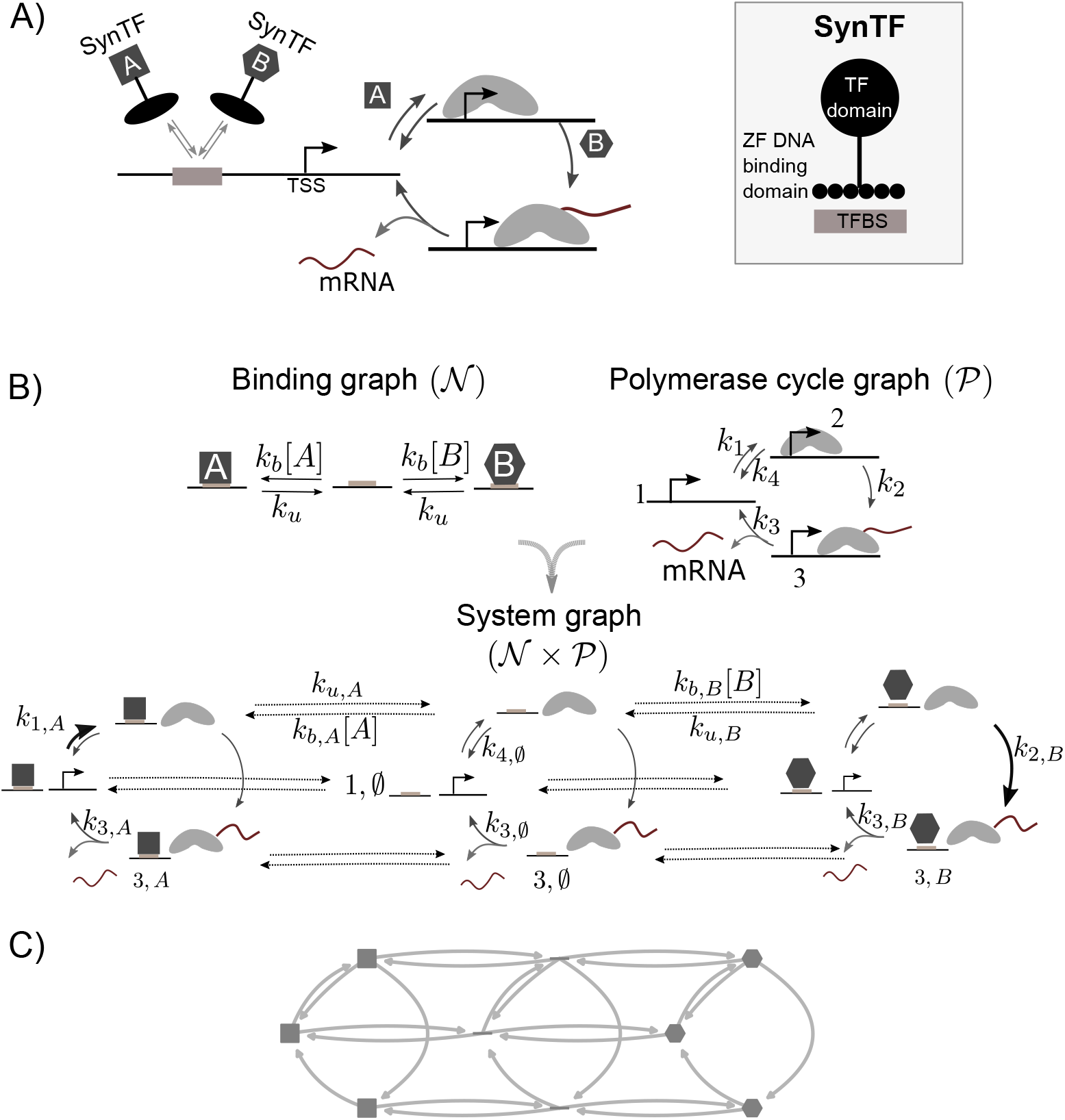
A model for kinetic synergy between two TFs sharing a site. A) Cartoon schematizing the strategy of this work to examine kinetic synergy: two synthetic TFs regulate a reporter (not shown) through a shared binding site. As an example, TF *A* controls the first step in the transcription cycle, and TF *B* controls the second step. B) Model used in this work. The graph product of a binding graph 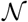, and a 3-state polymerase cycle graph 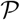, gives rise to the complete linear framework graph of the system 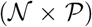. Only a subset of nodes and edges are labelled, for clarity. The horizontal edges from the central cycle to the outer cycles denote binding of each of the TFs, and the reverse edges denote unbinding. The three cycles allow us to account for the effect of the TFs, since the rates can be different depending upon the state of the binding site. As an example, the darker arrows denote the activator effect of *A* and *B* on the first and second transitions, respectively 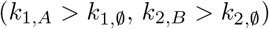. C) Schema of the full graph, as used in Figure 2.

## 4 Results

### 4.1 Mathematical model

We study how kinetic synergy emerges in a scenario where two TFs bind to a shared site in a regulatory sequence, such that only one TF can be specifically bound at any given time. Figure 1A schematizes this situation for a general 3-state transcription cycle, where TF *A* promotes the first step (illustrated as the assembly of the RNA polymerase complex), and TF *B* promotes a process downstream.

In order to model this system, we exploit the linear framework formalism, a graph-based approach to Markov Processes that can be used to model a diversity of biological processes in a biophysically realistic and mathematically tractable way (Gunawardena, 2012; Ahsendorf et al., 2014). We have previously applied this framework to study how binding interactions between TFs modulate gene expression by implicitly averaging over the states of the polymerase cycle (Estrada et al., 2016; Biddle et al., 2019; Park et al., 2019a). In contrast, in a previous study of kinetic synergy, we modelled the effect of TFs on a detailed transcription cycle but effectively combined their binding with their enzymatic effects (Scholes et al., 2017). Here we propose a model that unifies both approaches and doesn’t make assumptions about the binding reactions being on a different timescale than the polymerase cycle reactions, improving previous approaches in the literature (Li et al., 2018) (Methods, 6.1).

The system is represented by a graph (Figure 1B, 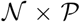), whose vertices are the biological states of interest, and the edges are the transitions between them, assumed to follow Markovian dynamics with infinitesimal transition rates corresponding to the graph edge labels. Structurally (i.e. ignoring edge labels) the graph for the complete system is the graph product between two simpler graphs: a binding graph and a polymerase cycle graph. The binding graph for our system of interest is represented in Figure 1B (Binding graph 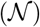), and consists of a binding site that can either be empty, bound by TF *A*, or bound by TF *B*. For the polymerase cycle (Figure 1B, Polymerase cycle graph 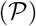), we consider the simplest cycle, with 3 states (labelled 1,2,3). The first transition is assumed to be reversible, and the other two irreversible in agreement with the macroscopic irreversibility of posttranslational modifications like phosphorylation, or the synthesis of mRNA. mRNA is assumed to be produced when the system transitions from state 3 to state 1. This simple graph can be interpreted in terms of empty transcription start site (TSS), assembled RNA polymerase, and C-terminal phosphorylated or elongating polymerase, although mapping onto specific states isn’t required to interpret the results. Given these two graphs, taking all pairwise combinations of their vertices (graph product) gives the complete graph 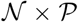 (Figure 1B).

TF binding on-rates (*k_b,X_, X* ∈ {*A*, *B*}, horizontal from the central cycle to the right and left) have dimensions of (concentration × time)^−1^, and binding off rates (*k_u,X_, X* ∈ {*A, B*}) have dimensions of (time)^−1^. The genomic context is modeled by the values of the basal rates over the polymerase cycle in the absence of TFs (central cycle). To incorporate the effect of a TF on a given transition, we assume that the TF only has effect while it is bound. The effect is then incorporated into the edge label (parameter value) for that transition, making it different for the cycle where the TF is bound than for the basal cycle. As an example, the darker arrows on the left and right cycles in Figure 1B, 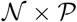, represent the activating effect of *A* and *B* on the first and second transitions, respectively. In this case, 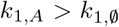, and 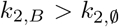. Similarly, repression could be included as well by a smaller value for a transition rate than the corresponding basal rate. For simplicity here we examine synergy between ”pure” activators only, defined by not decreasing the clockwise rates 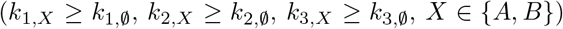 and not increasing the counterclockwise rate 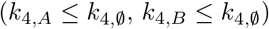.

We interpret the system in probabilistic terms, and assume each vertex of the graph holds the probability of the system being in that state. The transition rates then determine the time-evolution of the probabilities according to the Master Equation, which eventually reach a steady state (Methods, 6.1). Moreover, we assume first-order mRNA degradation. By taking the mRNA degradation rate as a constant that normalises the transition rates, the steady-state mRNA at a given concentration of *A* and *B* (*m*(*A, B*)*) is given by:

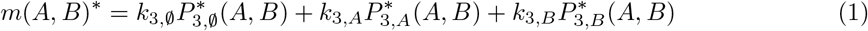

where 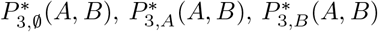 are the steady-state probabilities of state 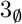, 3*_A_*, 3*_B_* at concentrations *A* and *B* of the TFs, and the rates are normalised by the mRNA degradation rate (Methods, 6.1). Given that we only consider the steady-state behaviour of the system, we use the same symbols to refer to the original rates and the normalised rates in order to avoid excessive notation. In the remainder of the paper, the rates will always be normalised.

The focus of the analysis is to compare this quantity when both TFs are present to that when only one is present and the other is at concentration 0 (synergy, below). Note that when only one or none of the TFs is present, *m*\* can be computed in the same way. In that case, the steady-state probabilities for those states corresponding to the absent TF being bound will be 0, and the rest will be redistributed according to the parameter values. The value of *m*\* in the absence of TFs (*m*\*(0,0)) corresponds to basal expression. For simplicity, the absence of a TF from the mathematical expressions below means it is at concentration 0.

### 4.2 A measure of synergy

Our interest is to understand how synergy emerges in this system. As shown by Scholes et al. (2017), if two TFs act on more than one step in the cycle, the overall effect may not be greater than additive even if they interact kinetically. This exemplifies that considering addition as a null expectation against which to define synergy, as has often been done in the literature, is model-specific. In order to provide a model-agnostic definition of synergy, here we consider a two-dimensional quantity that compares the steady-state expression when both TFs are present (*m*\*(*A, B*)) to the steady-state expression when either of them is alone, but at twice as much concentration (*m*\*(2*A*), *m*\*(2*B*)). In this way, the total concentration of TF is the same in the combined as in the individual situation. Enhanced expression in combination with respect to the strongest TF (the TF with a higher level of expression on its own), or reduced with respect to the weakest, must arise as a result of the functional interactions of the TFs over the cycle.

Positive synergy corresponds to higher expression in combination as compared to individually, and can be regarded as ”canonical” synergy in the sense of enhanced expression in combination: expression is greater than that of the strongest TF even if half the molecules are substituted by those of a weaker TF. We note however that the output does not have to be greater than additive to be considered positive synergy. Negative synergy corresponds to lower expression in combination, with expression lower than that of the weakest TF alone. Asymmetric synergy results when expression is increased only with respect to the weakest TF. In this case, it may be unclear whether there are any synergistic interactions. Potentially, these can still be detected depending on the extent to which the expression is reduced or increased with respect to the strongest or weakest TF, respectively. Thus, we propose to quantify synergy as a point in 2D, by comparing the effects of adding one TF to the other. This is quantified by *S_A,B_* (effect of *B* on *A*) and *S_B,A_* (the effect of *A* on *B*) as follows:

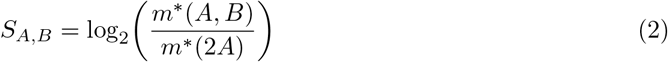

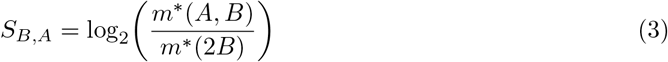

If *A* is taken to be the strongest TF, positive (green), asymmetric (blue) and negative (red) synergy map to 3 quadrants of a two-dimensional synergy space, as depicted in Figure 2A.

**Figure 2:**
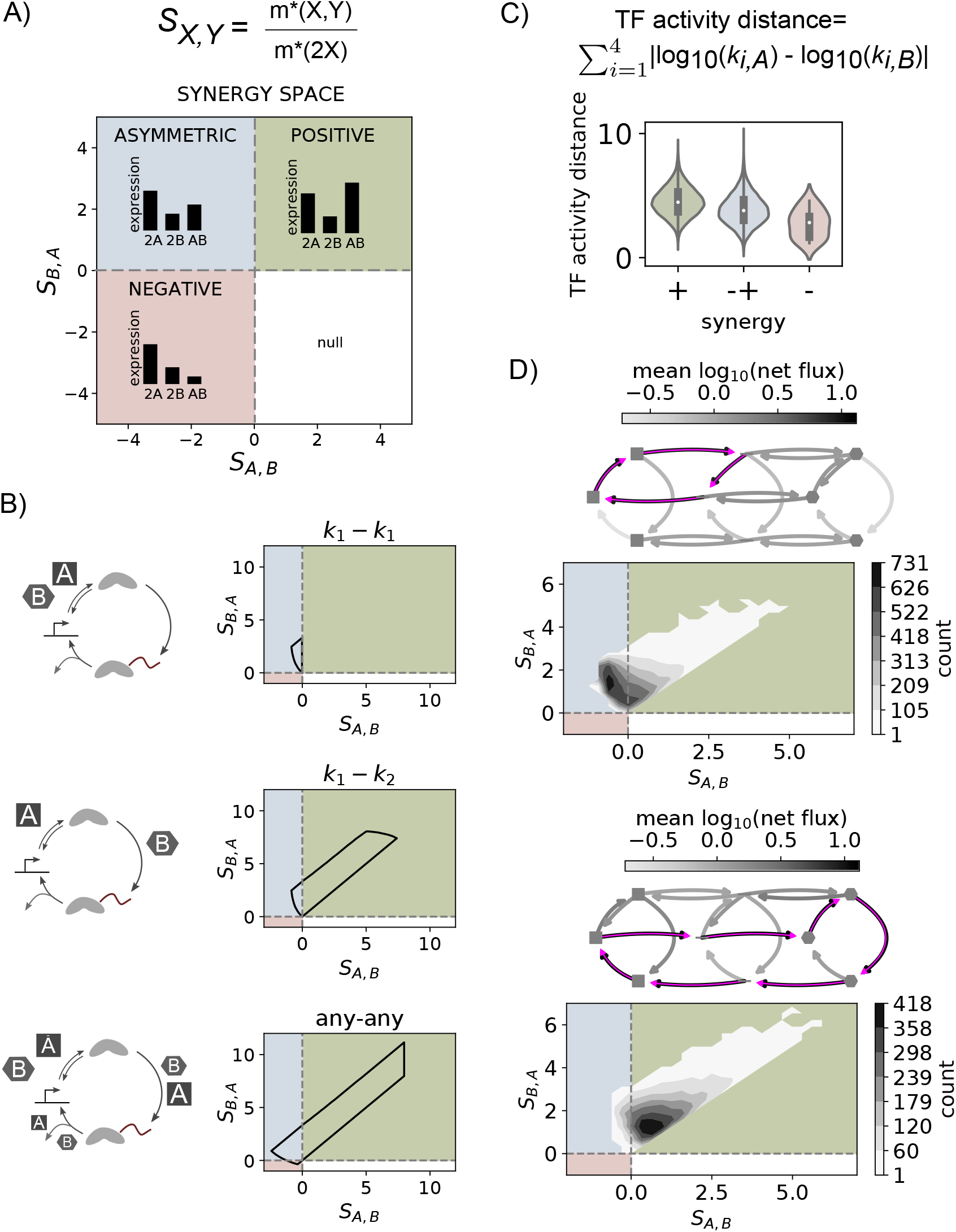
Positive, negative or asymmetric synergy emerge in the model depending upon parameter values. A) Synergy space. See text for details. B) Regions of the synergy space spanned by 3 regulatory strategies. Top: *A* and *B* act on the first step exclusively. Middle: *A* on one of the first two steps, *B* on the other one. Bottom: *A* and *B* act on any step (to various degrees). Constraints for the boundary search (Methods, 6.2, 6.3): parameter values between 1 and 10^4^, TF rates at most 1000 times larger than the basal rates for the clockwise (*k*_1_, *k*_2_, *k*_3_) or 0.001 times smaller for *k*_4_. Fold change in *m*\* for each TF individually (at concentration 2) with respect to the basal condition with no TF between 1 and 10. Figure S1B shows the results for more constrained parameters. C) Distribution of TF activity distances per synergy quadrant for a random sample of parameter sets under the same constraints as in the bottom panel in B (synergies are plotted as a scatterplot in Figure S1C). D) The two most prevalent dominant flux paths for the points used in the analysis in C. The arrow diagrams represent the model states and transitions, as schematized in Figure 1C; arrow greyscale intensity denotes the average probability net flux for that transition over all the parameter sets that share the dominant path highlighted in magenta. Note that reversible edges may appear in both directions if some parameter sets have net flux in one direction and others in the other. The distributions underneath show contours for the two-dimensional histogram of synergy values corresponding to those parameter sets that share the same dominant path. See also Figure S2.

### 4.3 Positive, negative or asymmetric synergy can theoretically emerge from two activators

We begin by exploring the theoretically possible synergistic behaviours between two activators 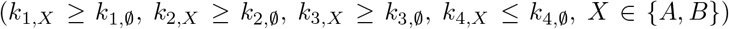. In line with our experimental system where TFs are comprised of the same binding domain (below), we assume that both TFs have the same binding kinetics (given by a binding rate *k_b_* and an unbinding rate *k_u_*) but different activation capabilities (given by the *k_i,X_*, *i* ∈ {1, 2, 3, 4}, *X* ∈ {*A, B*}). We assume the concentration unit is incorporated in the binding on-rates, such that both *A* and *B* are present at a concentration of 1 arbitrary unit each when they are both present together, and at concentration 2 when they are alone. In order to define the boundaries of the synergy space region that can be covered by the model under biologically-plausible parameter values and constraints (Methods, 6.2), we numerically sampled the parameter space using a biased sampling algorithm (Methods, 6.3). We explored the synergy space when TFs act on the same step, exclusively complementary steps, or all steps (Figure 2B).

As a control, we first explored the case where both TFs enhance the first step. Figure 2B-top shows that as expected, only asymmetric synergy appears in this case. Intuitively, if TF *A* drives stronger expression than TF *B* but both act on the same step, then mixing *A* with *B* can only reduce expression with respect to the strongest one, and increase it with respect to the weakest.

Next, we explored the case where TFs have complementary activities, with each TF enhancing either the first or second transition. Figure 2B-middle shows that this control strategy mostly results in positive synergy, but also covers a region of the asymmetric synergy quadrant (notice that the result is restricted to the upper diagonal region of the positive quadrant due to the definition of TF *A* as the strongest of the pair). A very similar result is obtained for any other pair of complementary rates (Figure S1A). The appearance of asymmetric synergy in this case shows that even if TFs have complementary activities, that may not be enough to enhance expression beyond that of the strongest TF when half of its concentration is substituted by the weaker TF.

TFs are often found to interact with a wide range of cofactors and regulators (Dingar et al., 2015; Kim et al., 2017; Carnesecchi et al., 2020), and it is therefore likely that they modulate multiple processes albeit with different strengths. Hence, we next considered a more general scenario where each TF can enhance any of the transitions to different extents (Figure 2B, bottom). In this case, a slightly higher region of the positive and asymmetric synergy quadrants are occupied, and slightly negative synergy can also emerge. We interpret this as an indication that under some parameter values, TFs can interfere with each other’s action and reduce the expression as compared to when only one of them is present.

For all these cases, the synergy space region that can be spanned by the model becomes smaller for more constrained parameter values, representing the assumption that the system has a smaller basal expression and TFs are weaker (Figure S1B).

### 4.4 The activity of the TFs over the cycle is not the only determinant of synergy

The original proposition of kinetic synergy stemmed from the assumption that synergy would emerge from TFs acting on different rate-limiting steps in transcription (Herschlag and Johnson, 1993). In the case of TFs with potentially overlapping effects, to what extent is positive synergy linked to TFs working exclusively, or nearly exclusively, on separate steps, so that they complement each other to enhance the cycle? In order to address this question, we looked at the correspondence between parameter values and synergy. For this, we generated a random sample of points that span a wide region of the synergy space (plotted in Figure S1C, Methods, 6.4). In order to quantify the degree of complementarity between the pair of TFs in a given parameter set, we use the following measure, which we call TF activity distance: the sum, over all the polymerase cycle transitions, of the absolute differences between the logarithms of the transition rates associated to each TF (Figure 2C). Similar TF parameter values result in a small distance value, whereas TFs with big differences in their rates, and therefore more divergent in their functions, result in a larger distance. As shown in Figure 2C, positive synergy tends to emerge at higher distances than asymmetric and negative synergies, suggesting more divergent functions is indeed linked to higher complementarity and thus higher positive synergy.

However, the distances that lead to asymmetric synergy and those that lead to positive synergy overlap, suggesting that the different functions of the TFs are not the only determinants of synergy output. When binning the distributions by the basal expression (steady state *m*\* in the absence of TFs) and binding and unbinding rates, these factors appear to be important as well: higher basal expression and higher binding and unbinding rates correlate with less distant TFs producing positive synergy (Figure S1D). In addition, the basal expression and binding rates also modulate the correlation between the distance of two TFs and the extent of positive synergy that they exhibit (Figure S1E).

Intuitively, for positive synergy to emerge, we would expect that each of the TFs binds and unbinds appropriately as to be able to exert its effect and not interfere with the binding and the effect of the other TF. In order to test the extent to which this is indeed linked to synergy, we looked at the steady-state probability fluxes in the graph. Given the irreversible nature of the transitions of the polymerase cycle, a net probability flux remains even when the system is at steady state. The flux of probability of the system is intimately linked to the production of mRNA, since mRNA is produced as the system transitions through the polymerase cycle. Formally, the flux from node *i* to node *j*, *J_ij_* is given by *J_i,j_* = *k_ij_ P_i_*, with *k_i,j_* the transition rate between *i* and *j*, and *P_i_* the probability of node *i*. In the case of irreversible edges, this equals the net flux. In the case of reversible edges, the net flux 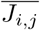 can be defined as 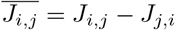, with *J_i,j_* > *J_j,i_*.

For the same sample of points (parameter sets) as in Figure 2C, we computed the net fluxes in the presence of *A* and *B*. Then, for each point, by starting at the polymerase-empty state with no TF bound (state 1, 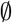 in Fig 1B, 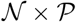) we followed the transition with a higher net flux, and repeated the same iteratively until reaching state 1, 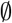 again or any other node already encountered. This generates what we call the dominant path of net fluxes over the graph. After computing the dominant path for each of the paramater sets, we quantified how many parameter sets share the same dominant path. For this analysis, we pulled together those pairs of paths that are mirror images of each other, since they are equivalent.

Out of all the parameter sets sampled, the majority correspond to one of either two paths, represented in Figure 2D. The most predominant involves the binding of one TF, transition over the first step (binding of polymerase), unbinding of the TF, and reversion to the empty state. The two-dimensional density plot below the flux diagram shows that the majority of the points with this dominant path of fluxes correspond to asymmetric synergy. In contrast, the second most frequent dominant path involves cycling over the whole graph, with the first two transitions occurring under one TF, and the last occurring under the other. In this case, the majority of the points are associated with positive synergy. The rest of the dominant paths that make up to 90% of all the dominant paths in the sample of points are shown in Figure S2. The density plots show that dominant paths are not uniquely associated to individual synergy classes, but there are clear biases, with positive synergy being mostly associated to dominant paths that traverse the whole graph, and asymmetric synergy linked to dominant paths that show nonproductive cycling. This agrees with the expectation that positive synergy should emerge when TFs act productively to enhance progression over the polymerase cycle, but also suggests that an intricate balance between all the transitions in the system is required for positive synergy to emerge.

### 4.5 Experimental evidence of kinetic synergy using a synthetic platform

The modelling results in the previous sections suggest that kinetic synergy can be observed from this single binding site circuit. In order to experimentally test this idea, we developed a reporter system in which synthetic TF fusions are recruited to a single binding site integrated into a mammalian HEK293 cell line (Methods, 6.6, 6.8) (Khalil et al., 2012; Park et al., 2019b; Israni et al., 2021). We selected five activation domains of mammalian TFs with a described diversity of functions in the literature. SP1 is a ubiquitous mammalian transcription factor whose mechanism of action has classically been linked to the recruitment of the transcriptional machinery (O'Connor et al., 2016). cMyc is also a ubiquitous regulator. It interacts with a diverse range of proteins, but its mechanism of action has been predominantly linked to processes downstream the recruitment of the transcriptional machinery, including pause-release (Rahl et al., 2010) and elongation via interaction with the elongation factor Spt5 (Baluapuri et al., 2019). BRD4 has also been described to have elongating activity, through the interaction with positive transcription elongation factor b (pTEF-b) (Yang et al., 2005; Moon et al., 2005). In addition, it has been involved in phase-separation at super-enhancers (Vasile et al., 2018), suggesting that BRD4 may also regulate other steps in the transcription cycle. Finally we chose the activation domain of HSF1, which has been described to have both initiating and pause-release stimulating activity, and a mutant version of it, which we call HSF1-m. This mutant was described to be elongation-deficient (Brown et al., 1998). Accordingly, these TFs can be broadly classified into either initiating (if they influence the recruitment of RNA polymerase) or elongating factors (if they influence a process downstream), as depicted in Figure 3A.

**Figure 3:**
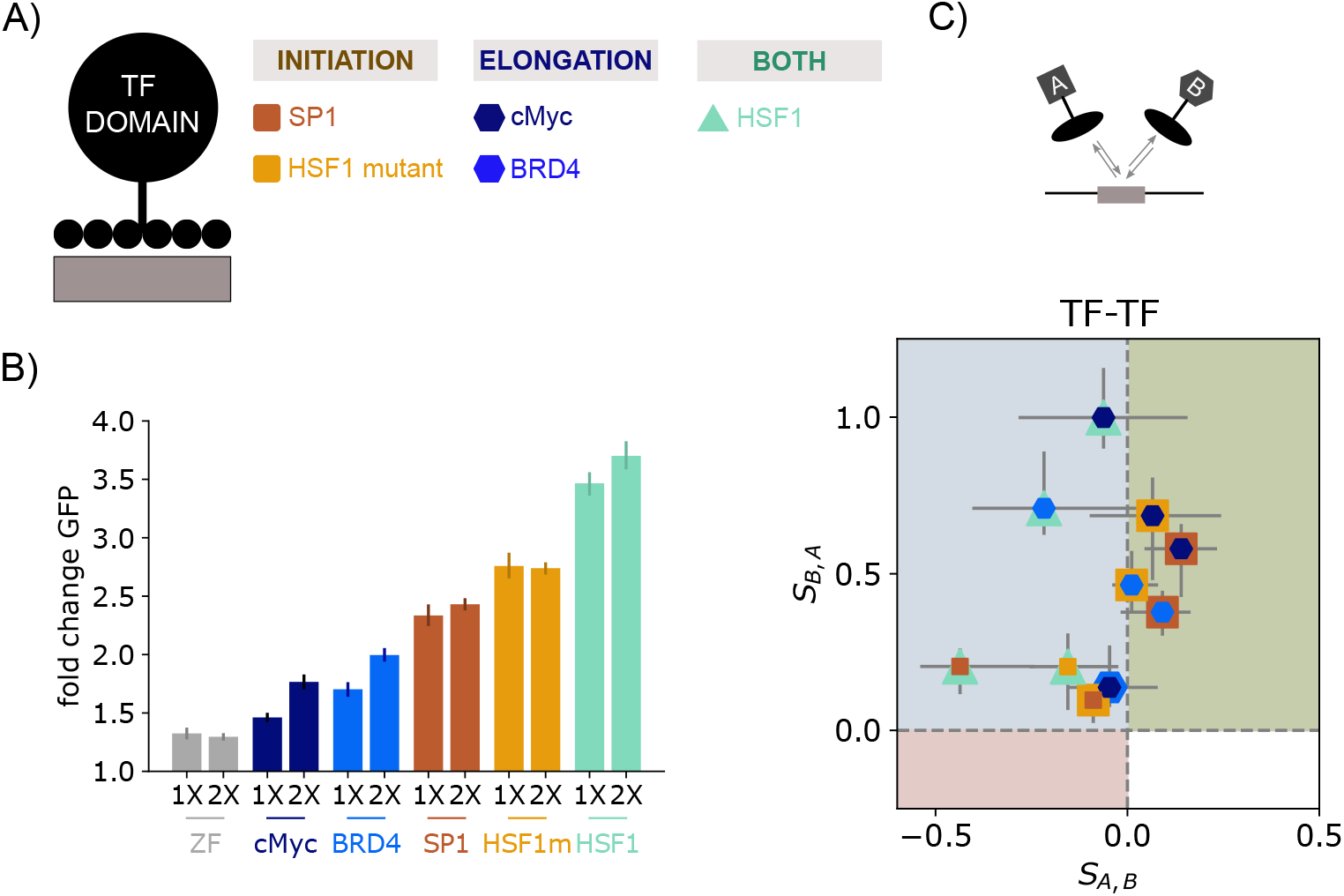
Experimental observation of kinetic synergy between 2 transcriptional activators on a synthetic platform. A) Schema of the synthetic TFs. B) Expression upon transfection with either 10 (1X) or 20 (2X) ng of one TF, or the ZF binding domain alone (grey). Error bars denote the 95% confidence interval for the mean GFP fold change, obtained from bootstrapping the mean GFP fold change values from all the experiments (biological replicates) for each condition. At least 3 biological replicates per condition, with 2-4 technical replicates each. C) Experimental synergy between two activators, defined as in Eqs. 2–3 (log_2_ of the ratio of average fold-change expression when 10 ng of each TF is transfected, over the average fold-change expression when 20 ng of one is transfected). TF *A* is the strongest of the pair in the single TF expression, as shown in the 2X conditions of panel B. Error bars denote ranges from at least three biological replicates, with 2-4 technical replicates each. Barplots corresponding to this data are shown in Figure S3F, and the synergy between each TF and the empty ZF is shown in Figure S3E.

We engineered synthetic TFs (synTFs) composed of an activation domain from the above-described TFs fused to a synthetic zinc finger (ZF) DNA binding domain (Methods, 6.6), designed to target a 20-bp binding site that does not natively exist in the mammalian genome sequence (Figure 1A, Figure 3A) (Khalil et al., 2012; Park et al., 2019b; Israni et al., 2021). This allows us to specifically recruit the activation domains to a reporter to assess their effects on transcription, while minimizing confounding effects from native TFs acting on the reporter. We then stably integrated into HEK293FT cells a reporter, composed of a single target binding site upstream of a minimal CMV (minCMV) promoter driving the expression of a destabilized EGFP (d2EGFP) (Methods, 6.8). Given its rapid turnover (Li et al., 1998), destabilized EGFP serves as a convenient genomic reporter of the mRNA expression level (Raj et al., 2006). The expression of the synTFs was induced by transient transfection of the synTFs, whose expression level can be controlled by the amount of the plasmids transfected (Figure S3A). We chose to transfect synTFs at either 10 or 20 ng to ensure that the concentration (i.e. expression level) of synTF is the limiting factor. Reporter expression outcome was assessed by quantifying GFP fluorescence using flow cytometry 48 hours later (Methods, 6.9, 6.10).

Figure 3B shows reporter activation by each of the synTFs. We observed similar activation strengths varying from about 1.5 fold change in GFP fluorescence to 4 fold change, with slight increases upon doubling the amount of TF transfected for most TFs. Such fold change up-regulation is in the range of physiological induction in mammalian signalling pathways (Strasen et al., 2018; Wong et al., 2019; Friedrich et al., 2019). A similar dose-dependent increase in reporter signal is also observed at the mRNA level (Figure S3B), supporting the use of GFP fluorescence to report on mRNA.

In order to assess the extent of synergy between pairs of TFs, we compared the fold-change in GFP fluorescence when TFs were transfected in pairs at 10 ng each, to that when only one is transfected at 20 ng. We used quantitative immunofluorescence targeting the HA-tag of the synTFs to verify that transient transfection of 20 ng of coding plasmids for a single synTF results in a similar synTF abundance distribution as when transfecting two TFs in combination at 10 ng each, despite some variability inherent to the transfection procedure (Figure S3C,D) (Methods, 6.13). Under these conditions, Figure 3C shows that both positive and asymmetric synergy appears (See Figure S3F for details). Consistent with the correlation in the model between TF activity distance and synergy class, the pairs exhibiting positive synergy (Fig 3C, green quadrant) correspond to those where each TF predominantly has been described to have either initiating or elongating factor activities. No TF was capable of increasing the expression from that driven by HSF1, which is the strongest synTF in the set and is described to have both initiating and elongating activities (Brown et al., 1998). However, different TFs reduced its expression to different extents, suggesting some functional interactions are occurring (e.g. compare the *S_A,B_* coordinate for SP1-HSF1 and cMyc-HSF1 in Figure 3C). For the pairs of TFs described to predominantly act upon the same step, almost no synergy was detected (SP1-HSF1m, cMyc-BRD4).

Figure 3B shows a very modest activation effect from the ZF alone (no TF activation domain) case. However, the combination with a full synTF only leads to asymmetric synergy (Figure S3E), with all TFs except HSF1 being reduced by the same extent, and HSF1 being reduced even further. This suggests that although the ZF may have a small effect perhaps by increasing the ability of the basal transcriptional machinery to bind, the positive synergy observed between pairs of TFs is most likely due to their activation domains, since the ZF only reduces expression when mixed with any of the TFs.

These results show that positive synergy can emerge experimentally even when the TFs share the same binding site. However, the effects are small. One potential reason is that the TFs are weak, in agreement with the model (Figure S1B). Moreover, the distributions of Figure S1D-E and the analysis of the dominant flux paths in Figure 2 point to binding and unbinding kinetics as important contributors to synergy as well. We now focus on this point.

### 4.6 Kinetic synergy depends upon the binding and unbinding kinetics

We explored how the synergy exhibited by a pair of TFs changes in the model as a function of either the unbinding or the binding rate. We began by examining the effect of the unbinding rate. To this end, we randomly sampled parameter sets for the basal rates over the polymerase cycle 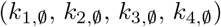 and binding and unbinding (*k_b_*, *k_u_*). For each of these basal sets, we sampled parameter values for pairs of TFs (*k*_1,*A*_, *k*_2,*A*_, *k*_3,*A*_, *k*_4,*A*_, *k*_1,*B*_, *k*_2,*B*_, *k*_3,*B*_, *k*_4,*B*_). For each pair, we varied the unbinding rate *k_u_* over a 2 order magnitude range, 10 fold up and down the basal value, and tracked the corresponding behavior over the synergy space. Given that the unbinding rate changes expression from each TF alone, we only considered those parameter sets where the strongest TF is the same across the unbinding rates considered, so that synergy is consistently defined throughout. Further details of this procedure are given in Methods, 6.5.

To classify the behavior over the synergy space systematically, we considered that the binding and unbinding rate are related to affinity by *K_a_* = *k_b_*[*TF*]/*k_u_*, and we used the relationship between changes in synergy and affinity so that the same criteria can be used to analyse the results when perturbing either the binding or the unbinding rate. We focused on the positive and asymmetric synergy behaviors, and used a 4-bit string that captures the behaviour at the affinity extremes: the first position denotes if *S_A,B_* is positive (p) or negative (n) at highest affinity, and the second position denotes the sign at the lowest affinity. The third and fourth positions denote whether *S_A,B_* and *S_B,A_* increase (i) or decrease (d), respectively. We disregard those situations where there is no change. As a result, there are theoretically 12 possible behaviors. We found that for some basal sets of parameters, changing the unbinding rate could result in all 12 possible behaviors, depending on the pair of TF parameter values. One such example is shown in Figure S4A, and selected examples are shown in Figure 4A. Similar results were found when modulating the binding on-rate *k_b_* (Figure S4B), which can be interpreted as modulating the baseline concentration of the TFs at 1X concentration.

**Figure 4:**
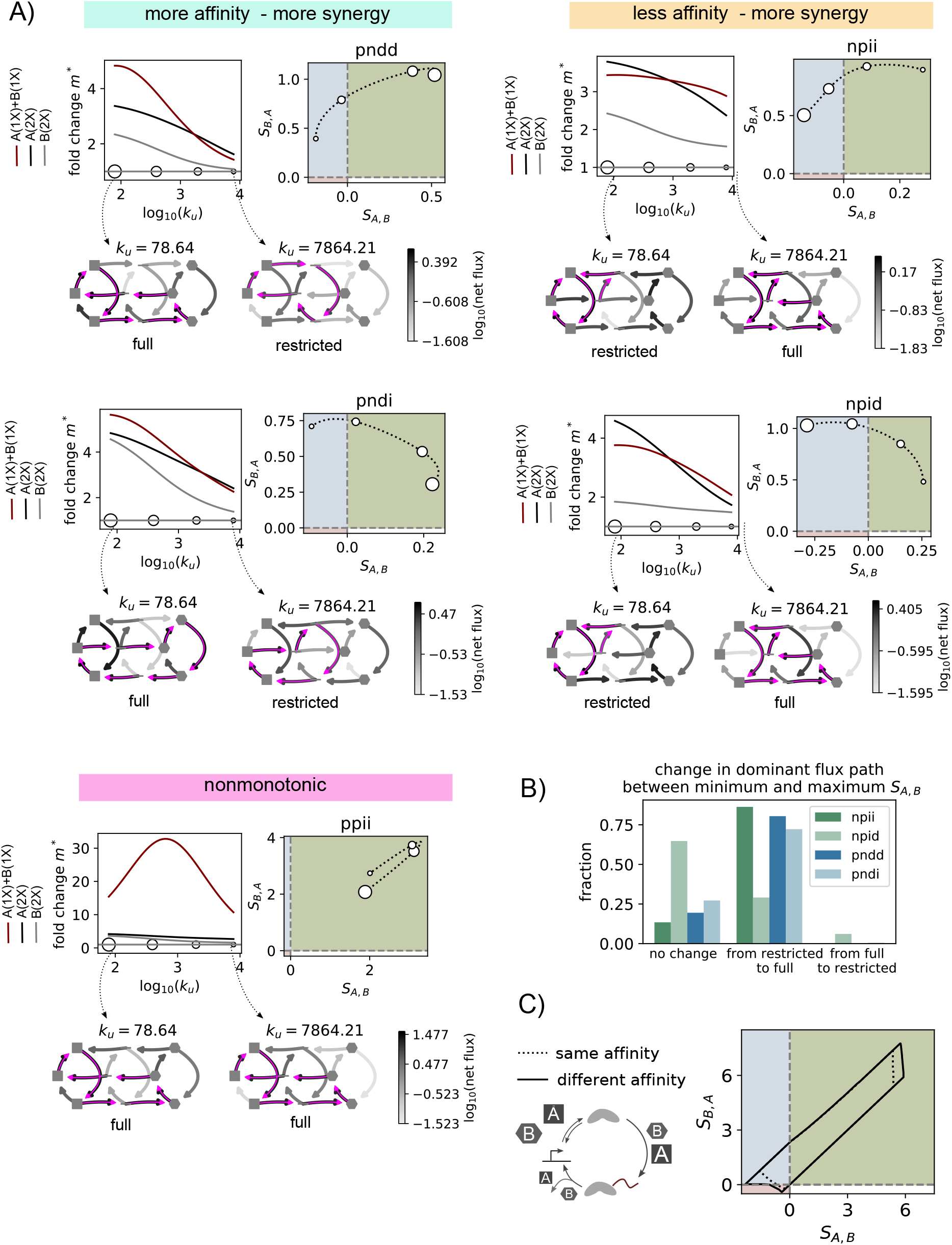
Synergy between a pair of TFs depends upon the binding and unbinding kinetics. A) Model examples for 5 sets of parameter values demonstrating the diversity in how synergy changes as a function of the unbinding rate. For each example, the top-left plot shows the fold change in expression as compared to no TF present, for each of the TFs at concentration 2 (black, gray), or both TFs at concentration 1 (maroon), as a function of the unbinding rate. The top-right plot shows the corresponding behavior in synergy space. The circles on the bottom of the top-left plot and those on the top-right plot correspond to the same values of synergy. Marker size is related to binding affinity (smallest marker: smallest affinity, highest unbinding rate). Shown below are the diagrams depicting the net fluxes (grey colormap) and dominant flux path (magenta) for the two extreme *k_u_* values. All examples share the same basal parameter values: 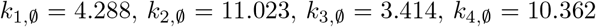. *k_b_* = 180.19. TF associated parameter values are as follows: pndd : *k*_1,*A*_ = 120.985, *k*_2,*A*_ = 154.358, *k*_3,*A*_ = 4.561, *k*_4,*A*_ = 2.854, *k*_1,*B*_ = 5.007, *k*_2,*B*_ = 25.685, *k*_3,*B*_ = 15.086, *k*_4,*B*_ = 2.083; pndi : *k*_1,*A*_ = 6.317, *k*_2,*A*_ = 517.659, *k*_3,*A*_ = 1433.877, *k*_4,*A*_ = 1.095, *k*_1,*B*_ = 11.275, *k*_2,*B*_ = 326.127, *k*_3,*B*_ = 15.328, *k*_4,*B*_ = 10.223; npii : *k*_1,*A*_ = 4.844, *k*_2,*A*_ = 6345.641, *k*_3,*A*_ = 151.500, *k*_4,*A*_ = 7.354, *k*_1,*B*_ = 4.504, *k*_2,*B*_ = 17.664, *k*_3,*B*_ = 2601.429, *k*_4,*B*_ = 3.088; npid : *k*_1,*A*_ = 6.784, *k*_2,*A*_ = 740.850, *k*_3,*A*_ = 56.436, *k*_4,*A*_ = 2.010, *k*_1,*B*_ = 4.821, *k*_2,*B*_ = 11.997, *k*_3,*B*_ = 909.506, *k*_4,*B*_ = 8.354; ppii : *k*_1,*A*_ = 937.265, *k*_2,*A*_ = 8084.904, *k*_3,*A*_ = 5.392, *k*_4,*A*_ = 1.982, *k*_1,*B*_ = 9.945, *k*_2,*B*_ = 18.372, *k*_3,*B*_ = 2047.513, *k*_4,*B*_ = 8.447; See also Figure S4. B) Quantification of the change in dominant path in the presence of both TFs, from the smallest to the largest *S_A,B_*. The parameter values were obtained from a rejection based sampling algorithm, as explained in section 6.5. The number of parameter sets analysed for each class are as follows: npii: 13103 parameter sets, corresponding to 214 basal parameter sets. npid: 4461, corresponding to 264 basal parameter sets. pndd: 2833, corresponding to 87 basal parameter sets. pndi: 2215, corresponding to 132 basal parameter sets. C) Region of the synergy space spanned by the model under parameter constraints determining weak basal expression and weak TFs: basal expression parameter values between 1-100 for clockwise rates, 100-1000 for 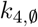. TF parameter values at most 100X greater (0.01X smaller for *k*_4_). Fold change from each TF alone at 2X concentration limited to 5. *k_b_* and *k_u_* are either same for both TFs (dotted line), or different (solid line). For the case of same binding, it is the same result as the dotted line in Figure S1B, right.

As expected from typical occupancy-based hypotheses, we found instances where increasing affinity led to an increase in synergy (Figure 4A, more affinity-more synergy), changing from asymmetric to positive. In contrast, we also found examples where even if the expression from the individual and combined TFs decreases with less affinity, synergy increases and can change from asymmetric to positive as affinity is reduced (Figure 4A, less affinity-more synergy). As seen in Figure S4 and depicted at the bottom of Figure 4A, we found many instances of nonmonotonic behaviour, where synergy was maximal at intermediate affinities.

To examine the relationship between the change in synergy class and the cycling over the system promoted by the TFs, we determined the dominant paths of net fluxes at steady state for parameter sets where synergy changes between asymmetric and positive or vice-versa as a function of the unbinding rate. We calculated the dominant path for the lowest and highest *S_A,B_* in the presence of both TFs. For each dominant path, we assessed whether it spanned nodes in each of the three binding configurations of the system (”full path”) or not (”restricted path”), as depicted for the corresponding examples of Figure 4A. Then, for each parameter set, we assessed whether the path type changed between the smallest and largest *S_A,B_* value, and plotted the quantification in Figure 4B. As expected, and in line with the examples in Figure 4A, the barplot shows that in the majority of the cases, the change from smallest to highest *S_A,B_* value correlates with a transition from a restricted to a full dominant path. For the case where increasing the unbinding rate causes synergy to increase only with respect to TF A (npid), we found many instances with no change of path class, and a small set where the relationship was reversed. This result aligns well with those of the previous sections, which show that the synergy of a pair of TFs ultimately depends on the overall system behaviour and the intricate balance between all the transitions. However, a major contributor to the synergistic behaviour of the TFs is the productive cycling over the system, with each TF binding and unbinding appropriately to allow the other to exert its effect. Therefore, we hypothesized that by combining pairs with different activation domains and affinities, synergy might be further enhanced. In agreement with this, the synergy space covered by the model expands slightly when weak TFs have different binding and unbinding rates, as compared to when their binding parameters are the same as we have considered in the previous analyses (Figure 4C). This suggests a scenario where the combinatorial effect of TFs can be flexibly tuned by the combined effect of their biochemical activities and binding.

## 5 Discussion

In eukaryotic transcription, combinatorial control occurs at multiple scales, with many TFs binding to a given enhancer, and many enhancers controlling the activity of a gene (Spitz and Furlong, 2012). Here we have focused on the first scale, and have investigated how synergy between TFs can emerge as a result of the kinetics of the system. Though kinetic synergy was theoretically proposed almost 30 years ago (Herschlag and Johnson, 1993), its experimental demonstration has been challenging, largely due to the confound of cooperative binding interactions. To circumvent this limitation, we have focused on a scenario where only one TF can be specifically bound at any given time. By forcing the TFs to act separately in time, their functional interactions can be revealed. In order to reason about this scenario, we have proposed a minimal biophysical model that explicitly accounts for the kinetics of the binding as well as the functional effects of the transcription factors over the transcription cycle. The model reveals that synergy between a pair of TFs is not an intrinsic feature of the pair, but depends upon the balance between their binding and their functional effects. This work gives yet another example of the power of synthetic biology to answer fundamental biological questions (Crocker et al., 2017; Park et al., 2019a; Bashor et al., 2019).

### A measure of synergy

In order to quantify synergy, it has been common to measure the deviation from additivity, under the assumption that if TFs do not interact, then their combined effect should be the sum of the effects obtained when each TF is present alone (Carey, 1998). Multiplicativity has also been taken as a measure of synergy (Bintu et al., 2005a). However, in Scholes et al. (2017) we showed that when TFs interact functionally on a 2-step cycle, additivity or multiplicativity is only expected under very restricted circumstances. In order to provide a model-agnostic measure of synergy, here we propose to compare expression when both TFs are present together, to expression when only one of them is present, under the same total TF concentration. By having the same total TF concentration in both cases, changes in the expression when there are two TFs as compared to only one must be due to their functional interactions, and therefore provides evidence of synergy. In addition to positive synergy, we define asymmetric and negative synergy. This enables the quantitative characterization of the synergy between a pair of TFs as a function of a variable of the system, by looking at the corresponding trajectory in synergy space. Although this measure is particularly suited for the single binding site scenario explored here, we suggest it could also be used to quantitatively characterize the response to combinations of TFs in a more natural scenario where each TF binds to distinct sites.

### A model that explicitly accounts for the interplay between TF binding and polymerase activity

In order to reason about the single binding site experiment, we have developed a model with details of both the binding of the TFs and the progression over the polymerase cycle. This model brings together the two main modelling frameworks of transcription in the literature, where either the binding is taken implicitly (e.g. Scholes et al., 2017), or the polymerase cycle is not detailed (e.g. Estrada et al., 2016). In contrast to other attempts in the literature (Li et al., 2018), we don’t make assumptions about the timescales of the binding and unbinding of the TFs with respect to those of the biochemical transitions over the polymerase cycle. This provides greater generality. In addition, the model can easily be extended to include more polymerase states and more binding sites for other TFs or coregulators, if such details become relevant in future studies. One of the simplifying assumptions of the model is that TFs only exert their effect while they are bound. We note that this doesn’t necessarily have to be the case, since they may act through other cofactors that can remain bound even if the TF unbinds. This could be easily incorporated at the expense of more states and parameters. However, we think it wouldn’t fundamentally change our conclusions, since there would also be an interplay between the binding kinetics of these other components and the kinetic effects on the cycle.

We have explored the behavior of the model in parameter space under the assumption that the system is at steady state. This is a widely used assumption and reasonable for our experimental setup, given the time between transfection and measurement of mRNA levels. However, one of the contexts where combinatorial control is most relevant is development, and many developmental processes may be too fast to allow for a steady state to be reached. In this case, it may become important to explicitly incorporate the time delay that emerges from polymerase travelling along the gene body, which we have not accounted for. Although at steady state this is likely to be effectively incorporated by the parameter of the last transition rate in the polymerase cycle, it could have important implications when considering how synergy emerges in transient regimes, and will be a relevant point to consider in future studies.

### Kinetic synergy can emerge when two TFs time-share a binding site

We have found that extensive positive synergy is theoretically possible in the case where two activators bind to the same site on DNA. Our analysis shows that this is due to TFs productively enhancing the polymerase cycle when acting in combination, by binding and unbinding appropriately to allow each TF to exert its effect. We note that the extent of positive synergy experimentally observed is small compared to the regions covered by the model. In the model, we have found that the region of the synergy space is reduced as more constraints on the parameters are imposed, especially when constraining the extent to which a TF can enhance a given rate, and the expression fold change that it causes. Therefore, the small synergy experimentally observed suggests that the synTFs have relatively weak effects, in agreement with the small fold-change activation that they produce.

According to the model, synergy between a pair of TFs is strongly influenced by their binding kinetics. Theoretically, both the binding on-rate and off-rate can modulate the synergy exhibited by a pair of TFs, and lower affinity can increase the synergy observed for a pair of TFs, even if this reduces expression from the TFs acting individually. In some cases, the compromise is evidenced as a nonmonotonic effect of affinity upon synergy. ZFs with different binding affinities can be obtained by introducing mutations in the ZF scaffold that are known to mediate non-specific interactions with DNA (Khalil et al., 2012). In future work, synTF variants can be used to systematically explore the role of binding affinity on synergy. Given the small effects of individual synTFs on transcription, which may be weakened further by affinity mutations, it will be critical to have fine control over synTF expression, and dynamic measurements in single cells are likely to be informative.

Our previous analyses had suggested that assessing synergy might be a way to elucidate the mechanism of action of TFs (Scholes et al., 2017). However, the current analysis shows that this is confounded by the effect of the binding kinetics. Moreover, parameter constraints that generate positive synergy in the model also generate asymmetric synergy. In this case, even if TFs may have complementary activities, their binding patterns may be imbalanced and may not allow productive interaction. In the case where either of the TFs works exclusively on one of two complementary steps, this contrasts with the finding of exclusively greater-than-additive behaviour by Scholes et al. (2017), highlighting the importance to account for the binding kinetics.

The model also shows that when TFs have overlapping activities, negative synergy can emerge even if individually they act to enhance the cycle. Again, this arises due to an imbalance between the timescales of their binding and functional effects, where in combination they interfere with each other. However, the extent of this effect is small and requires very fine tuned parameter sets, as evidenced by the low numbers of points in this region obtained from pure random sampling. In agreement with this, we did not robustly observe negative synergy experimentally.

### Implications for gene regulation in natural scenarios

In endogenous enhancers, some TFs do have overlapping binding sites as in our setup (Han et al., 1998; Pan and Nussinov, 2011; Cheng et al., 2013). However, most typically, each TF has its own binding site. Even in this case, binding kinetics may still be important. The residence time of the TF on the DNA must be long enough for it to be able to exert its function. However, it is plausible that there could be interference either directly or through recruited cofactors, such that output may be maximized at intermediate affinities. This could be another reason behind the widespread presence of relatively low-affinity binding sites in eukaryotes (Ramos and Barolo, 2013; Farley et al., 2016; Crocker et al., 2016; Kribelbauer et al., 2019), and the observation of fast TF binding kinetics (Paakinaho et al., 2017; Li et al., 2019; Donovan et al., 2019). Moreover, tuning binding site affinity might be an effective way to modulate expression beyond fully adding or removing a binding site, which could have evolutionary implications (Kurafeiski et al., 2019). Along the same lines, kinetic synergy relaxes the need for strict arrangements between binding sites, another typical feature of eukaryotic transcriptional control (Kulkarni and Arnosti, 2003; Junion et al., 2012; Smith et al., 2013).

TF activity has often been considered to be modular. In this view, the activity of the activation domain is independent of that of the binding domain, which is assumed to be important only to target the TF to specific sites on the genome (Ptashne, 1988). Evidence against this model includes allosteric interactions between the DNA binding domain and the activation domain (Li et al., 2017), and the observation that the activation domain may be involved in DNA recognition (Brodsky et al., 2020). Adding to this, our work highlights the importance of considering TFs as a unit, where the binding and activation domains together dictate the effect of the TF. Our study emphasizes the value of considering an integrated view of transcriptional control, where the effect of a TF has to be understood in terms of the other components of the system.

## 6 Methods

### 6.1 Modelling details and the linear framework

In this work we have used the linear framework formalism to model the interplay between the TF binding and their effects on the transcription cycle. This framework was introduced in (Gunawardena, 2012) and we have previously exploited it to study other problems in gene regulation. Ahsendorf et al. (2014); Estrada et al. (2016); Biddle et al. (2019, 2021) can be consulted for details. We outline the main features here.

A biological system is represented by a finite, directed, labelled graph *G* with labelled edges and no self-loops. The graph represents a coarse-grained version of the system of interest, with the nodes being the states of interest, and the edges the transitions between them. The edge labels are the infinitesimal transition rates for the underlying Markov process, with dimensions of (time)^−1^, and they include terms that specify the interactions between the graph and the surrounding environment. For example, the transitions that represent the binding of a TF have edge labels that include the TF concentration, which is assumed to remain constant over time (i.e. TF is sufficiently in excess that, to a good approximation, binding does not reduce the concentration of free TF available for binding).

The graph defines the time-evolution of the probability for each state of the system (vertex) as follows. Assume that each edge is a chemical reaction that follows mass-action kinetics with the edge label as the rate. Since each edge has only one source vertex, the resulting dynamics is linear and is described by a matrix equation,

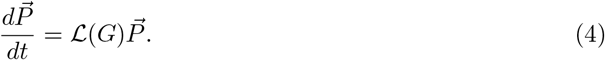

Here, 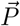 is the column vector of state probabilities at time *t*, with dimension *n*, and 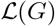 is the Laplacian matrix of the graph. Eq. 4 is the master equation, or Kolmogorov forward equation, of the underlying Markov process.

For a strongly connected graph, the system has a unique steady state, where 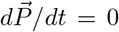. The steady-state probability values for each state are computed by summing over the products of the rate labels for each of the spanning trees rooted at that state, and normalising appropriately (see Estrada et al. (2016) for details).

The mRNA concentration *m* is assumed to evolve according to:

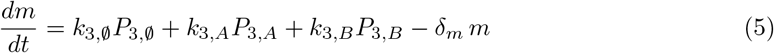

where the *P*_3,*X*_ are the probabilities of states 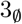, 3*_A_*, 3*_B_* at a given time (Figure 1B). By assuming steady state, setting *dm/dt* = 0, and dividing by *δ_m_*, we obtain the expression for the steady state mRNA (* denotes steady state):

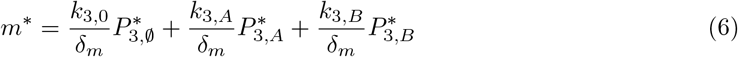

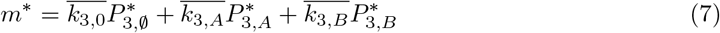

This gives Eq. 1 of the main text, where the overbars are dropped for simplicity. In the parameter exploration, we directly sample on the normalised rates.

### 6.2 Biologically plausible ranges for parameter values

We considered a biologically plausible range for the normalised parameter values to be between 1 and 10^4^, according to the following reasoning:

The events from the binding of the polymerase complex until the production of an mRNA molecule involve many biochemical reactions, including the binding interactions associated with the assembly of the pre-initiation complex, the phosphorylation of the C-terminal domain of RNA polymerase and other post-translational modifications (Schröder et al., 2013), as well as the biochemistry associated to elongation. Our 3-state cycle is therefore a coarse-grained representation of all these processes. In order to determine biologically plausible parameter ranges, we searched for measurements of reaction rates for these processes, and normalised those to typical rates of mRNA degradation, taken to have typical half-lives between 10 min (0.00116 *s*^−1^) and 5 h (3.85 × 10^−5^ *s*^−1^) (Sharova et al., 2009; Chan et al., 2018).

For a reaction at a rate of 0.7 s^−1^ (~1 s half-life), normalizing by the mRNA degradation rates would result into a normalised range of 600-18000.

For a rate of 0.07 *s*^−1^ (~10 s half-life), the normalised range would be 60-1800.

For a rate of 0.016 *s*^−1^ (~1 min half-life), the normalised range would be 10-300.

And for a rate of 0.00116 *s*^−1^ (~10 min half-life) the normalised range would be 1-30.

These values are consistent with measurements of various transcription-associated biochemical reactions: the *in vitro* rate of pre-initiation complex assembly was found to vary over ranges on the order 10^−3^ s^−1^ (Ferguson et al., 2001) to 0.1 *s*^−1^ (Kugel and Goodrich, 1998), and the rate of promoter opening/escape was reported to be 0.002 *s*^−1^ (Kugel and Goodrich, 1998). Pause stability is estimated to be from 3 s to 20 min (Wissink et al., 2019). And the TF residence time can be from just a few seconds to a few minutes (Paakinaho et al., 2017; Mehta et al., 2018).

Therefore, we took a range of 1-10^4^ for our parameter values. We note that we also checked the results with smaller ranges, around slower rates for the polymerase cycle, but found that it didn’t affect the results qualitatively, only reduced the synergy region as shown in Figure S1B.

### 6.3 Synergy space boundary for a regulatory strategy

In order to determine the region of the synergy space that can be spanned by a given regulatory strategy, we used a biased random sampling algorithm, modified from that in Estrada et al. (2016). Parameters were chosen from a given range of normalised rate values, and TFs were assumed to at most modify the basal rates by a certain factor (see figure captions for the values corresponding to each figure). A maximum fold change for expression in the presence of one TF alone (at 2X concentration) was also pre-specified, such that parameter sets that generate expression outside this range were discarded. The steps of the algorithm are as follows:

1. Define constraints and two-dimensional grid of synergy values. Initialize with the hyperparameters (below).
2. Randomly sample parameter values from their range (in log scale) until 10 points are found that fall in different cells of the grid.
3. Until convergence: at each iteration, search the surrounding parameter space of each boundary point (see below) and keep the new parameter sets that generate synergy values not already found (empty cells). Convergence is determined by 3000 iterations where no new points occupying empty cells are found.

In order to search the surrounding parameter space of a given parameter set (point in synergy space), we followed 3 strategies (each point was modified using the 3 strategies at each iteration, provided there were sufficient points for steps 2 (10) and 3 (100)):

1. Randomly select a few parameter values and modify them.
2. ”Pull” towards a target point in the direction determined by the centroid and the point being modified, away from the boundary: for 500 trials or until convergence, slightly modify the parameter set, and keep the new one if it generates a point in synergy space closer to the target.
3. ”Pull” in the direction (approximately) perpendicular to the tangent between the point being modified and its neighbor, as in 2.

The algorithm depends on various hyperparameters: probability of selecting a parameter value for mutation (0.2, 0.5), probability of replacing an already-existing boundary parameter (0.2,0.6), width of the interval around a parameter value to sample for new parameter values (in log (base 10) scale: [−2,2],[−1.5,1.5],[−1,1]). Searches were run for all 12 combinations of hyperparameters, and results were merged together.

The boundary search code is available at https://github.com/rosamc/GeneRegulatoryFunctions. The rest of the code to reproduce the calculations and figures in the paper is available at https://github.com/rosamc/kinsyn-2021.

### 6.4 Random sample of points in synergy space

In order to randomly sample parameter values in the synergy space we followed a rejection sampling approach. Parameters were sampled logarithmically from its predefined range (1-10^4^). The constraints on the maximum fold change effect on the polymerase cycle rates by the TFs were checked, as well as the constraint on the expression fold change by each of the TFs at 2X. We collected 1 million parameter sets that satisfied the constraints. Then, in order to have a more uniform distribution of points over the synergy space, we binned the synergy space into a grid with bins every 0.025 *S_A,B_* and *S_B,A_*, and kept one parameter set per bin. We repeated the procedure 10 times. The resulting points are those in Figure S1C.

### 6.5 Exploration of synergy as a function of binding or unbinding rate

In order to explore how synergy depends upon the binding and unbinding rates, we generated sets of basal parameters by randomly sampling on a logarithmic scale the basal rates between 1 and 10^4^, and the binding and unbinding rates between 10^1.5^ and 10^3^. For each of these basal parameter sets, we generated parameter sets corresponding to the TF-associated parameters, and we kept 1000 such parameter sets that satisfy the following constraints: i) TF-associated parameter values at most 1000X the respective basal ones (0.001X for counterclockwise rate *k*_4_); ii) fold change in expression by each TF individually at 2X concentration between 1 and 5; iii) TF A is consistently the strongest of the pair when the binding or unbinding rate is changed by a factor *f*, where *f* spans 10 logarithmically spaced values between 0.1 and 10. For each parameter set that satisfied the constraints, we determined the class of behaviour in synergy space as a function of the change in the binding or unbinding rate over this two-order magnitude range, and saved for downstream analysis those parameter sets where the absolute value of the change in both *S_A,B_* and *S_B,A_* was at least 0.05.

### 6.6 Construct design and cloning

The reporter construct consists of a single synthetic zinc finger binding site (CGGCGTAGCCGATGTCGCGC) upstream of a minimal CMV promoter (taggcgtgtacggtgggaggcctatataagcagagctcgtttagtgaaccgtcagatcgcctgga) driving d2EGFP (EGFP destabilized with signal peptide for fast degradation (fusion with aa 422-461 of mouse ornithine decarboxylase)).

synTF fusion proteins containing an activation domain of interest fused to an N-terminal zinc-finger binding domain with a GGGGS flexible linker were driven under control of a ubiquitin promoter and contain a 5’ sv40 nuclear localization sequence, C-terminal HA and rabbit globin polyA 3’ UTR. Genome-orthogonal zinc fingers were previously developed to target 20-bp sequences that minimize identity with the reference human genome (Israni et al., 2021; Park et al., 2019b). The following protein domains were selected and conjugated as respective activation domains according to previous studies:

SP1 (Residues 263 – 499) [PMID: 8278363] NITLLPVNSVSAATLTPSSQAVTISSSGSQESGSQPVTSGTTISSASLVSSQASSSSFFTNANSYSTTTTTSNMGIMNFTTSGSSGTNSQGQTPQRVSGLQGSDALNIQQNQTSGGSLQAGQQKEGEQNQQTQQQQILIQPQLVQGGQALQALQAAPLSGQTFTTQAISQETLQNLQLQAVPNSGPIIIRTPTVGPNGQVSWQTLQLQNLQVQNPQAQTITLAPMQGVSLGQTSSSN

cMyc (Residues 1-70) [PMID: 12177005] MDFFRVVENQQPPATMPLNVSFTNRNYDLDYDSVQPYFYCDEEENFYQQQQQSELQPPAPSEDIWKKFEL

BRD4 (Residues 1308-1362) [PMID: 24860166] PQAQSSQPQSMLDQQRELARKREQERRRREAMAATIDMNFQSDLLSIFEENLF

HSF1 (Residues 370-529) [PMID: 9606196] PEKCLSVACLDKNELSDHLDAMDSNLDNLQTMLSSHGFSVDTSALLDLFSPSVTVPDMSLPDLDSSLASIQELLSPQEPPRPPEAENSSPDSGKQLVHYTAQPLFLLDPGSVDTGSNDLPVLFELGEGSYFSEGDGFAEDPTISLLTGSEPPKAKDPTVS

HSF1 mutant (Residues 370-529, F418A, F492A, F500A) [PMID: 9606196] PEKCLSVACLDKNELSDHLDAMDSNLDNLQTMLSSHGFSVDTSALLDLASPSVTVPDMSLPDLDSSLASIQELLSPQEPPRPPEAENSSPDSGKQLVHYTAQPLFLLDPGSVDTGSNDLPVLAELGEGSYASEGDGFAEDPTISLLTGSEPPKAKDPTVS

### 6.7 Cell culture

HEK293FT cells (Thermo Fisher Scientific) were used as a background cell line in this study. Cells were cultured in DMEM with L-glutamine, 4.5g/L Glucose and Sodium Pyruvate (Thermo Fisher Scientific) supplemented with 10% FBS (Clontech), GlutaMAX supplement (Thermo Fisher Scientific), MEM Non-Essential Amino Acids solution (Thermo Fisher Scientific) and 1% penicillin-streptomycin (Thermo Fisher Scientific). Cells were maintained at 37°C with 5% CO2 in a humidified incubator, with splitting every 2-3 days.

### 6.8 Genomic integration of reporter constructs

Reporter lines were generated by site-specific integration of reporter constructs into HEK293FT cells using CRISPR/Cas9 mediated homologous recombination into the AAVS1 (PPP1R2C) locus as previously described (Park et al., 2019b). Briefly, 60,000 cells were plated in a 48-well plate and transfected the following day by PEI with a mixture of the following: 70ng of gRNA_AAVS1-T2 plasmid (Addgene 41820), 70 ng of VP12 humanSpCas9-Hf1 plasmid (Addgene 72247), and 175 ng of donor reporter plasmid. Donor reporter plasmids contain flanking arms homologous to the AAVS1 locus, a puromycin resistance cassette, and constitutive mCherry expression. After transfection, cells were cultured in 2 mg/mL puromycin selection for at least 2 weeks with splitting 1:10 every 3 days. Monoclonal populations for reporter cell lines were isolated by sorting single cells from this population into a 96-well plate and growing cell lines from each well. A minimum of 6 monoclonal cell lines that express high level of mCherry protein were transiently transfected with a strong synTF activator (HSF1 or VP16) and a monoclonal cell line to be used going forward was selected based on the fold-change of GFP expression relative to basal GFP level.

### 6.9 Transient transfection

Stable reporter cell lines were transfected with synTF plasmid constructs using polyethylenimine (PEI, Polysciences) as described in (Park et al., 2019b). 60,000-100,000 cells/well were plated in 48-well plates and transfected the following day with a total of 10ng per synTF, unless otherwise noted, with single stranded filler DNA (Thermo Fisher Scientific) up to 200ng total. 50ng of pCAG-iRFP720 (Addgene, #89687) was used as a transfection control plasmid. Two days after transfection, cells were collected and prepared for flow cytometry, unless otherwise noted.

### 6.10 Flow cytometry and data analysis

For each measurement, cells were harvested and run on an Attune NxT (Thermo Fisher Scientific) or LSR II (BD) Flow Cytometer equipped with a high-throughput auto-sampler. A minimum of 10,000 events were collected for each well and were gated by forward and side scatter for live cells and single cells, as described in (Park et al., 2019b). Cells were then gated by iRFP for transfection-positive populations. The geometric mean of GFP fluorescence distribution was calculated in FlowJo (Treestar Software). GFP expression fold-change was determined by normalizing with mean GFP intensity of the reporter only control. Flow cytometer laser/filter configurations used in this study were: EGFP (488 nm, 510/10), mCherry (561 nm, 615/25), iRFP-720 (638 nm, 720/30). All flow cytometry measurements were performed in technical replicates. Considering together all replicates from all experiments with the same transfection condition, we checked for consistency and discarded technical errors. This removed the cMyc 2X condition in one of the experiments since it yielded an aberrantly low fold change. Moreover, we removed 4 additional replicates, each from a different condition, that had a fold change that was above/below two standard deviations from the mean considering all replicates for that condition together.

### 6.11 Western blotting

A reporter cell line was transfected with indicated amounts of ZF-HSF1 (0, 10, 20, 50, 100, 150, 200ng) in a 48-well plate at a cell density of 1×105 per well. After 2 days, cells were rinsed with PBS and lysed with 200 *μ*L of NuPAGE LDS sample buffer (Thermo Fisher Scientific), followed by 5 seconds of sonication. Whole cell lysates were mixed with NuPAGE Sample Reducing agent (10X, Thermo Fisher Scientific) at 95°C for 5 minutes. Samples were then loaded into a 4-12% NuPAGE Bis-Tris Mini Protein precast gel (Thermo Fisher Scientific) and were run at 200V for 30 minutes in NuPAGE MES SDS Running Buffer. Separated proteins were transferred to a PVDF membrane using P0 protocol of iBlot2 system (Thermo Fisher Scientific). Membranes were blocked for 1hr at room temperature in blocking solution (5% w/v nonfat dry milk in 1X PBST) with gentle rocking. The membranes were probed with anti-HA (1:4000; Abcam ab9110) and anti-GAPDH (1:1000; Abcam ab9485) antibodies at room temperature for 1 hour with gentle rocking. The membranes were washed in PBST three times for 5 minutes each, and incubated with a goat anti-rabbit IgG-HRP antibody (1:2000; Abcam ab6721). The target proteins were visualized by chemiluminescence using SuperSignal West Pico PLUS substrate (Thermo Fisher Scientific) and an iBright Western Blot Imaging Systems (Thermo Fisher Scientific). Quantification of band intensities was carried out using FIJI (Schindelin et al., 2012).

### 6.12 Quantitative Real-Time PCR

1 × 10^5^ Hek293 reporter cells were seeded one day prior to transfection in 6cm culture dishes. Transfection was performed with the indicated amounts of synTF plasmid as described above for flow cytometry experiments using polyethylenimine (PEI) (polyscience) or Lipofectamine 3000 (Thermo Fisher Scientific). Two days post transfected, cell pellets were harvested and mRNA was extracted using the RNeasy Mini Kit (Qiagen). 500 ng extracted total RNA was reverse transcribed into cDNA for each sample. Reverse transcription was performed using Protoscript II reverse transcriptase (New England Biolabs) and oligo-dT primers (New England Biolabs). Quantitative real-time PCR was performed in triplicates using iTaq™Universal SYBR®Green reagent (Bio-Rad) on a CFX96 PCR machine (Bio-Rad). Primers were used in a final concentration of 243.2 nM. *β*-actin expression was used as a reference gene for relative quantification of RNA levels. Used primer sequences are (5’-3’):

Actin_fwd: GGCACCCAGCACAATGAAGATCAA;
Actin_rev: TAGAAGCATTTGCGGTGGACGATG;
eGFP_fwd: AAGTTCATCTGCACCACCG;
eGFP_rev: TCCCTTGAAGAAGATGGTGCG;

### 6.13 Comparison of synTF distribution across transfection conditions using quantitative immunofluorescence

#### 6.13.1 Immunostaining

0.5 × 10^5^ cells were seeded on poly-Lysine coated high-precision glass coverslips (18 mm round, #1.5) in 12-well culture plates one day prior to transfection. Transfection was performed as described for flow cytometry experiments. A total amount of 200 ng DNA (20 ng synTFs and 180 ng ssDNA) was used for transfection experiments. PEI was scaled to 12-well plate volume of 100 *μ*L total transfection mix. 48 h post transfection, cells were washed with 1x PBS, fixed with 2% PFA (Fisher Scientific) and blocked for 30 min with 10% Goat serum (VWR) in 1x PBS after washing. Immunodetection was performed with HA-tag (6E2) mouse antibody (Cell Signaling) 1:1000 in 1%BSA/PBS overnight. Cell were washed with 0.1% Triton X-100 and incubated with anti-mouse IgG Alexa Fluor 488 (#4408, Cell Signaling) antibody 1:1000 in 1%BSA/PBS for 1h. After washing with 0.1% Triton X-100, nuclei were stained with 2 *μ*g/mL Hoechst-33342 (Thermo Fisher Scientific) and mounted on glass slides using Prolong Gold Antifade (Thermo Fisher Scientific). Image acquisition was performed at least 16 h after mounting slides.

#### 6.13.2 Fluorescence microscopy

Images were acquired as single-plane multipoint positions on a Nikon Ti2 inverted microscope upon illumination by a Lumencor Sola 395 Light Engine and a Plan Apo VC 20x objective (NA 0.75). The following filter sets were used. Alexa Fluor 488: excitation FF01-466/40, emission FF03-525/50, dichroic FF495-Di03 (all Semrock); Hoechst-33342: excitation ET395/25x, emission ET460/50m, dichroic ET425lp (all Chroma). Detection was performed with a Hamamatsu ORCA Flash 4.0 LT camera. NIS elements software for image acquisition was used.

#### 6.13.3 Image Processing

Images were extracted from nd2 files, separated as .tif-files per channel and field of view. CellProfiler 4.0 (McQuin et al., 2018) was used for image segmentation and measuring nuclear fluorescence intensity. A pipeline was customized based on the pipeline for Human cells provided by the CellProfiler project. Nuclei segmentation was performed based on Hoechst-33342 staining using Otsu thresholding and a nuclear diameter range of 15 - 50 pixels. Objects outside that range and touching the border of images were excluded. Touching objects were distinguished based on fluorescence intensity and object intensity was calculated for the segmented nuclear area in all channels.

#### 6.13.4 Data analysis

The integrated fluorescence intensity (FI) calculated per nucleus for anti-HA-488 staining (detecting HA-tagged synTF) was used for further data analysis using custom scripts in Matlab for data processing. In case of multiple datasets of the same condition, FI distributions were joined. Background fluorescence was defined based on nuclear fluorescence intensity in untransfected controls. Data was normalized to the 5th-95th percentile to remove outliers from imperfect segmentation due to clumping of nuclei. The median was calculated and a threshold of 2.5 fold of the median was determined to identify positive transfected cells (Figure S3C). Nuclei with FI above this threshold were considered as positive transfected with any synTF condition described (Figure S3D). Distributions are plotted as probability density functions (PDF) using ksdensity function in Matlab. The 5th-95th percentile of values above threshold for each condition was taken to remove outliers for each condition to compare the distributions of synTF abundance in each dataset (Figure S3C).

## 7 Acknowledgements

We thank members of the DePace, Khalil and Gunawardena labs for helpful discussions. Fluorescence Microscopy experiments were performed at The Nikon Imaging Center at Harvard Medical School. We thank Jennifer Waters and Rylie Walsh for guidance and support. This work was supported by NSF grant MCB-1713855 (A.S.K.)/ MCB-1715184 (A.H.D), NIH grant 1R01GM122928 (A.H.D. and J.G.), NIH grant R01EB029483 (A.S.K.), NSF CAREER IOS-1452557 (A.H.D.), EMBO Fellowship 683-2019 (R.M.C.) and the Giovanni Armenise Harvard Foundation (A.H.D.). A.S.K. also acknowledges funding from the NIH Director’s New Innovator award (1DP2AI131083-01), an NSF CAREER award (MCB-1350949), and a DARPA Young Faculty award (D16AP00142). A.H.D. also gratefully acknowledges funding from the Lynch Foundation Systems Biology Fellowship, the McKenzie Family Charitable Trust Systems Biology Fellowship Fund, and the John and Virginia Kaneb Fellowship award at Harvard Medical School.

## 8 Supplementary Figures

**Figure S1.**
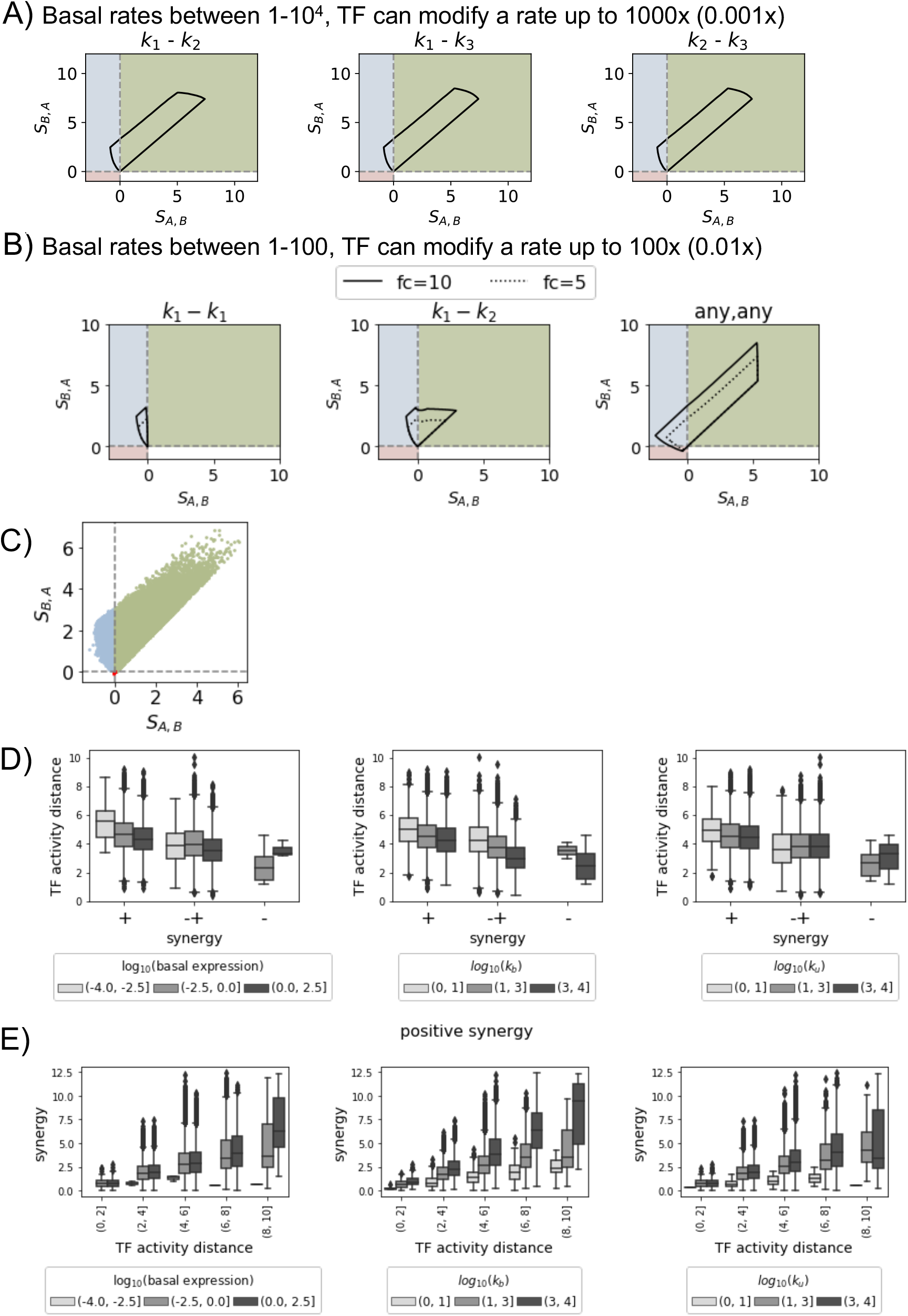
Characterization of the model behavior in synergy space. A) Region of the synergy space spanned by the model when each TF acts uniquely on one of two steps, indicated in the title of each plot. The first plot is the same as in Figure 2B middle. Parameter values in the range between 1 and 10^4^. Parameters for the TFs at most 1000 times larger than the basal parameters for the clockwise rates (*k*_1_, *k*_2_, *k*_3_) or 0.001 times smaller for *k*_4_. Fold change in *m*\* for each TF individually with respect to the basal condition with no TF bound between 1 and 10. B) Region of the synergy space from each of the 3 regulatory strategies in Figure 2B for more constrained parameters, representing weaker TFs: parameter values for the transitions over polymerase states in the range between 1 and 100 for the clockwise rates, 100-10000 for 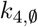; parameters for the TFs at most 100 times larger than the basal parameters for the clockwise rates (*k*_1_, *k*_2_, *k*_3_) or 0.01 times smaller for *k*_4_. Fold change in *m*\* for each TF individually with respect to the basal condition with no TF bound between 1 and 10 (solid line) or 1 and 5 (dashed line). C) Random sample of points where both TFs act on any step, randomly sampled under the same constraints as in Figure 2B bottom (Methods, 6.4) and used for the distributions in Figure 2C and panels D and E of this Figure. D) Distribution of TF activity distances as a function of synergy class, binned by basal expression (expression in the absence of TFs, left), binding on-rate (middle), binding off-rate (right), for the points in panel C. E) Distribution of synergy values as a function of TF activity distance and binned by basal expression, binding on-rate or binding off-rate for the points with positive synergy in panel C (green points).

**Figure S2.**
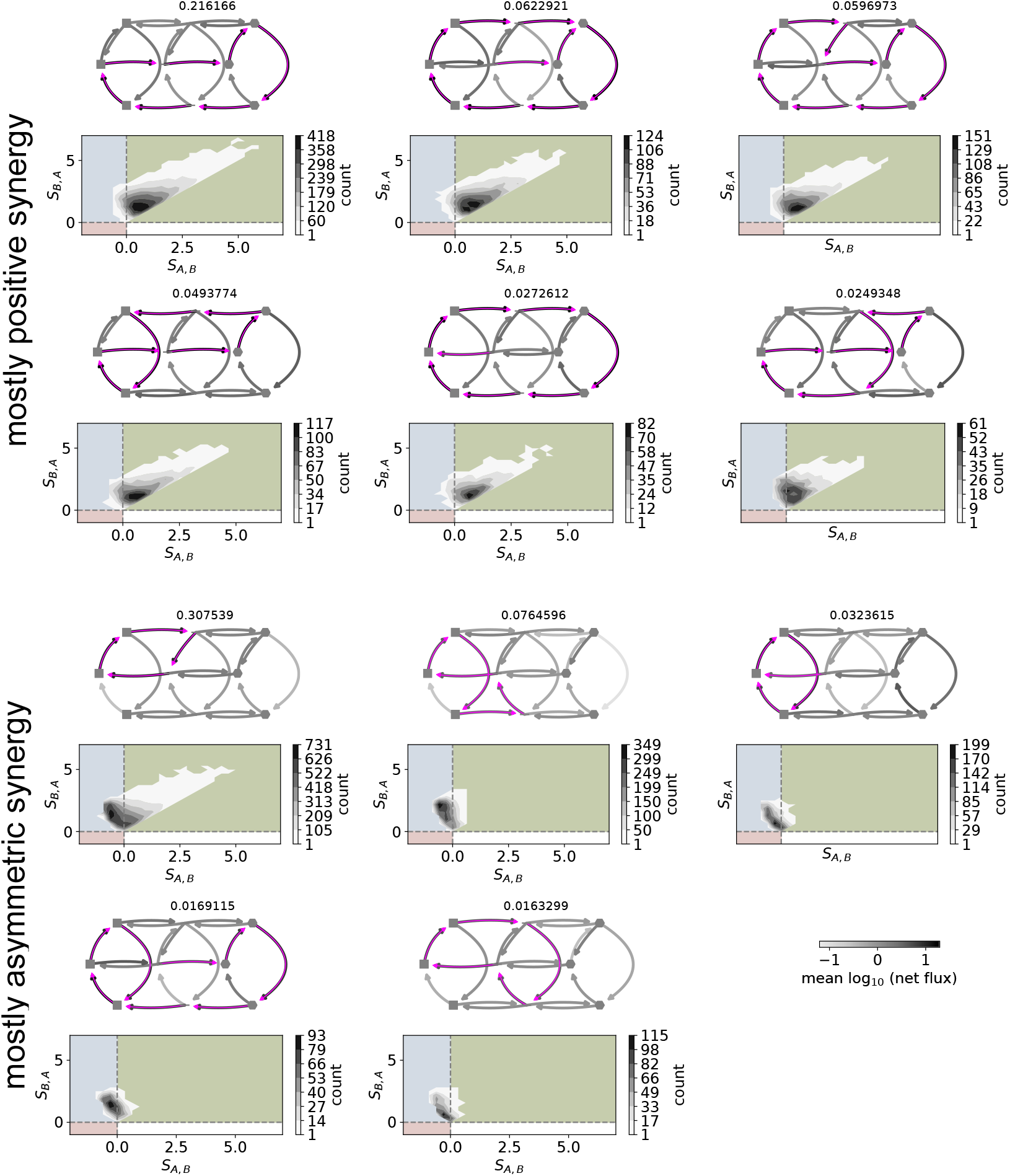
Top 90% net flux dominant paths for each parameter set in Figure S1C. For each path, the representation is as in Figure 2D. The top diagram shows the dominant path (magenta arrows) and the arrow greyscale shade encodes the average net flux for the transitions. The lower plot shows the distribution of synergy values that correspond to that dominant path. For each of the two groups, dominant paths are sorted according to their frequency, indicated on top of each. Note that the first path of each group corresponds to the plot in Figure 2D.

**Figure S3.**
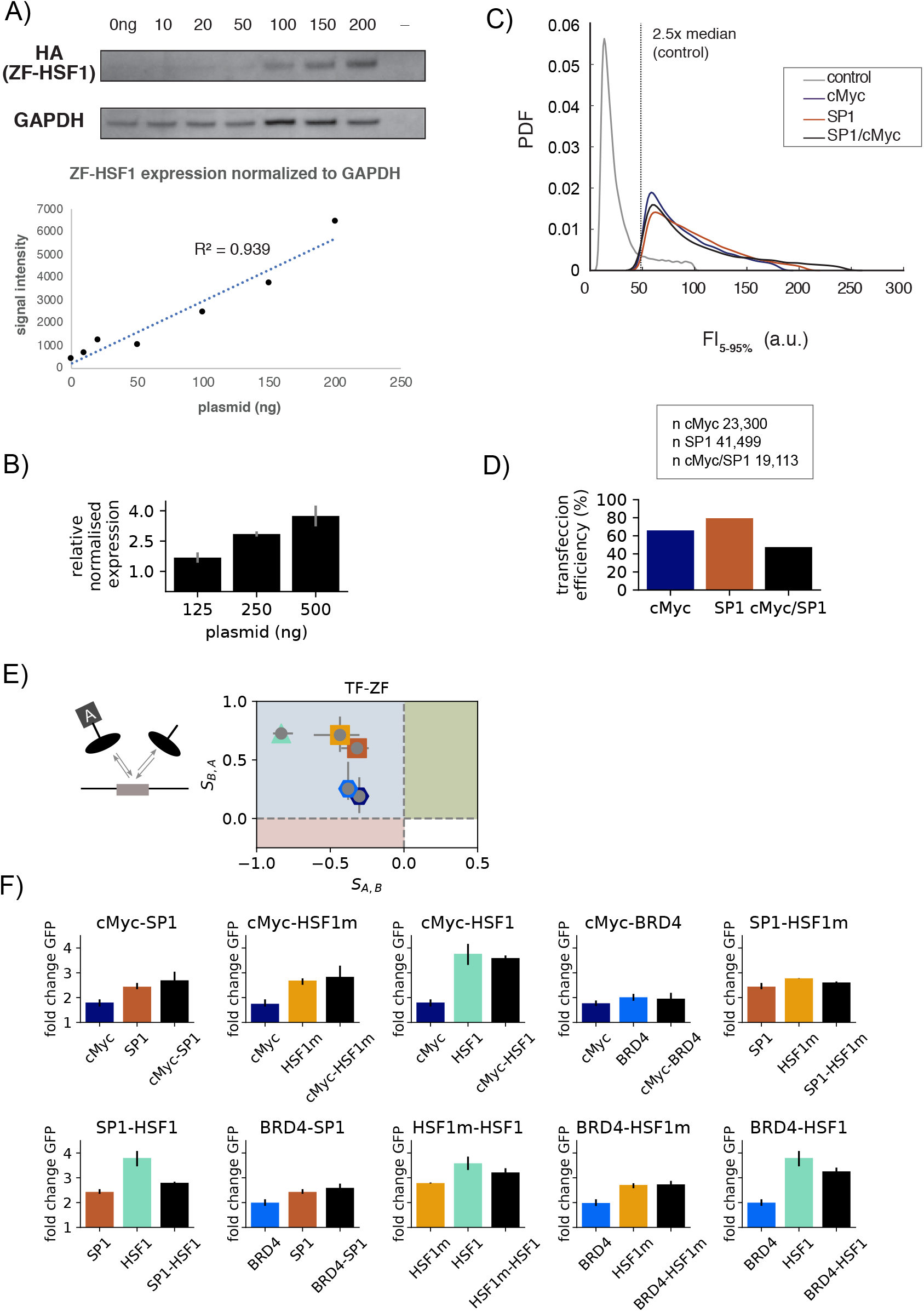
Experimental observation of synergy between a pair of TFs. A) Western Blot and corresponding quantification of the ZF-HSF1 protein as a function of ng of plasmid transfected (Methods, 6.11). B) Normalised GPF expression as measured by qPCR in response to SP1 synTF. The concentrations of plasmid used are scaled by the number of cells (Methods, 6.12) so that 125 and 250 ng are approximately equivalent to 10 and 20 ng in the flow cytometry experiments. Error bars denote SEM from technical replicates. C) Distribution of integrated Fluorescence intensity (FI) in arbitrary units (a.u.) as quantified from segmented nuclei for the control (gray, no synTF) and transfected samples using quantitative immunofluorescence targeting the HA-tag of synTFs. Data falling within the 5th-95th percentiles is shown for each dataset. Dashed line: background threshold defined as 2.5x median of control FI. SP1, c-Myc and SP1/c-Myc curves represent the 5th-95th percentile of values above threshold. (See Methods, 6.13 for details) D) Bargraph showing the percentage of positive transfected cells determined based on the background threshold shown in panel C. n is the number of quantified cells for each dataset above threshold. E) The combination of a synTF with the ZF alone only generates asymmetric synergy, where expression is between that of the ZF and that of the full TF. Error bars denote the ranges of the data. At least 3 biological replicates per combination, with 2-4 technical replicates each. F) Details of the fold change in expression for the conditions that generate the synergy plot in Figure 3C. Error bars denote the 95% confidence interval for the mean GFP fold change, obtained from bootstrapping the mean GFP fold change values from all the experiments for each condition. At least 3 biological replicates per combination, with 2-4 technical replicates each.

**Figure S4.**
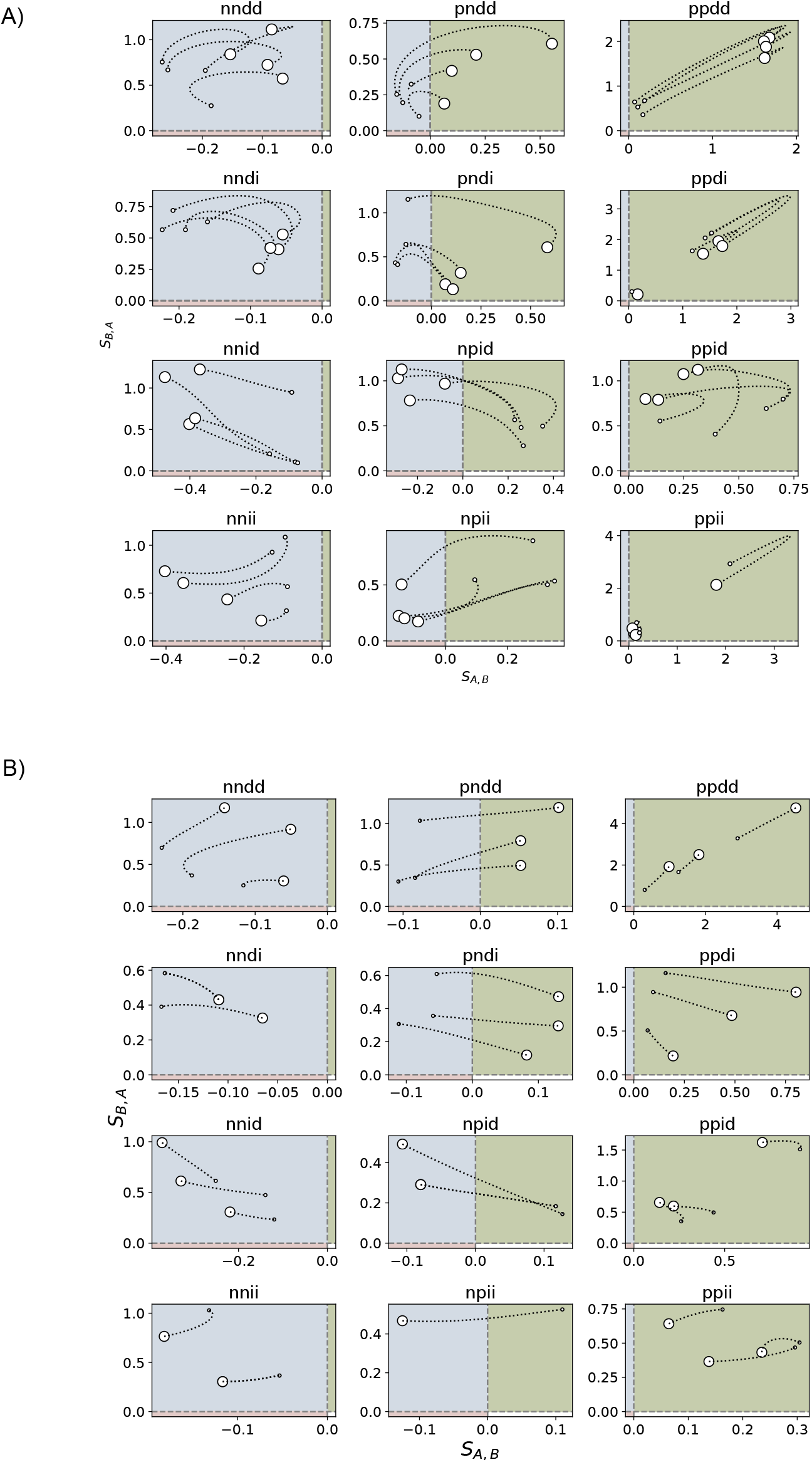
Synergy between a pair of TFs depends upon the binding and unbinding kinetics. A) Example of the 12 possible changes in synergy when the unbinding rate (*k_u_*) is modulated, for a common set of basal parameter values and different TFs. The biggest marker denotes highest affinity (lowest unbinding rate), and the smallest marker denotes lowest affinity (highest unbinding rate). The subplot titles indicate the behaviour as the unbinding rate is increased: both *S_A,B_* and *S_B,A_* decrease: nndd, ppdd, pndd. *S_A,B_* decreases but *S_B,A_* increases: nndi, pndi, ppdi. *S_A,B_* increases but *S_B,A_* decreases: nnid, npid, ppid. Both increase: nnii, npii, ppii. All lines share the same set of basal and binding/unbinding rates, but each corresponds to a given set of TF parameter values. Results have been selected out of all those found from a rejection-based sampling random search of parameter values. B) Example of the 12 possible changes in synergy when the binding rate (*k_b_*) is modulated. As in A, each line corresponds to a pair of TF parameter values, but all of them share the same basal and binding and unbinding rates. The biggest marker denotes highest affinity (largest binding rate), and the smallest marker denotes lowest affinity (lowest binding rate). See Methods 6.5 for details.

## Notes

### Competing Interest Statement

The authors have declared no competing interest.

